# Dynamic proteomics profiling of *Legionella pneumophila* infection unveils modulation of the host mitochondrial stress response pathway

**DOI:** 10.1101/2020.05.19.105395

**Authors:** Julia Noack, David Jimenez-Morales, Erica Stevenson, Tom Moss, Gwendolyn Jang, Nevan J. Krogan, Danielle L. Swaney, Shaeri Mukherjee

## Abstract

The human pathogen *Legionella pneumophila (L.p.)* secretes ~330 bacterial effector proteins into the host cell which interfere with numerous cellular pathways and often regulate host cell proteins through post-translational modifications. However, the cellular targets and functions of most *L.p.* effectors are not known. In order to obtain a global overview of potential targets of these effectors, we analyzed the host cell proteome, ubiquitinome, and phosphoproteome during *L.p.* infection. Our analysis reveals dramatic spatiotemporal changes in the host cell proteome that are dependent on the secretion of bacterial effectors. Strikingly, we show that *L.p.* substantially reshapes the mitochondrial proteome and modulates mitochondrial stress response pathways such as the mitochondrial unfolded protein response (UPR^mt^). To our knowledge, this is the first evidence of manipulation of the UPR^mt^ by a bacterial pathogen in mammalian cells. In addition, we have identified a previously uncharacterized *L.p.* effector that is targeted to host cell mitochondria and protects mitochondrial network integrity during mitochondrial stress.

## INTRODUCTION

Pathogenic bacteria commonly use their secretion systems to inject several proteins (effectors) into the host cell cytoplasm that manipulate and hijack cellular pathways. Studying these effectors have often revealed key insights into host cell biology (Bhogaraju et al., 2019; Cornejo et al., 2017; Kalayil et al., 2018; Mukherjee et al., 2011; Noack and Mukherjee, 2020; Qiu and Luo, 2017a). The intracellular bacterial pathogen, *Legionella pneumophila (L.p.),* is of special interest as it injects a remarkable number (> 330) of effector proteins via its Dot/Icm Type IV secretion system (T4SS) into the host cell (Finsel and Hilbi, 2015). These effectors rewire the host cell to avoid fusion of the *Legionella-containing* vacuole (LCV) with the endo-lysosomal compartment and establish a safe, endoplasmic reticulum (ER)-derived replicative niche (Cornejo et al., 2017). Several *L.p.* effectors regulate host cell proteins through post-translational modifications (PTMs) such as phosphorylation, ubiquitylation, and methylation (Michard and Doublet, 2015) and have led to the discovery of novel and fascinating biochemical mechanisms. For instance, *L.p.* studies have led to the discovery of completely novel PTMs, such as phosphocholination (Mukherjee et al., 2011), unique biochemical mechanisms, such as the non-canonical protein ubiquitination by bacterial ubiquitin ligases (Bhogaraju et al., 2016; Kalayil et al., 2018; Qiu et al., 2016; Wang et al., 2018), or novel regulatory roles of PTMs, such as translation inhibition through phosphorylation of Hsc70 by the *L.p.* kinase LegK4 (Moss et al., 2019). As the molecular targets and functions of most *L.p.* effectors are yet to be uncovered (Qiu and Luo, 2017b), studying the effects of these proteins on host cell pathways offers a great potential for the discovery of new cell biology.

A few targeted proteomics studies have been performed in order to identify novel host cell factors required for *L.p.* replication (Bruckert and Abu Kwaik, 2015; Ivanov and Roy, 2013; Schmölders et al., 2017; Urwyler et al., 2009). For instance, the ubiquitinome of *L.p.* infected cells revealed many interesting insights into the regulation of innate immune and mTOR signaling during infection (Ivanov and Roy, 2013). However, this approach relied on stable cell lines expressing tagged ubiquitin, a process that can presumably increase non-specific ubiquitination (Emmerich and Cohen, 2015; Peng et al., 2017). To our knowledge, a systematic time-dependent profiling of the phosphoproteome, ubiquitinome, and proteome has not yet been conducted. Therefore, we decided to perform a global analysis of the entire proteome, ubiquitinome and phosphoproteome during the course of infection with *L.p..* To distinguish between T4SS-dependent and -independent effects, we used the *L.p.* WT strain or the isogenic *ΔdotA* mutant, which lacks the T4SS and is cleared via the endosomal/lysosomal pathway. Our analysis provides a comprehensive resource which highlights T4SS-dependent, spatio-temporal changes in the host cell proteome during *L.p.* infection. Here, we identify the mitochondrial proteostasis network as a novel target of *L.p.* effectors. In response to infection with *L.p.,* the abundance, phosphorylation, and ubiquitination of several mitochondrial proteins involved in mitochondrial protein import, gene expression and protein folding were strongly regulated in a T4SS-dependent manner. The changes in the mitochondrial proteome show hallmarks of mitochondrial stress responses such as the mitochondrial unfolded protein response (UPR^mt^). The integrated stress response (ISR) pathway is known to be activated during the UPR^mt^ (Mottis et al., 2019). Interestingly, during *L.p.* infection, the transcription of mitochondrial stress genes was induced through an ISR-independent mechanism. Furthermore, *L.p.* infection suppressed the ISR, even upon exposure to a small molecule inducer of the UPR^mt^. As such, the translation of the ISR transcription factors ATF4 and CHOP was suppressed during *L.p.* infection. Instead, *L.p.* infection allowed the selective upregulation of the basic-region leucine zipper (bZIP) domain transcription factor ATF3 both at the level of transcription and translation. We believe that this is the first evidence of pathogenic UPR^mt^ activation that is independent of the ISR pathway. Furthermore, we show that the i-AAA mitochondrial protease YME1L1 contributes to the reshaping of the mitochondrial proteome upon *L.p.* infection by degrading the inner mitochondrial membrane (IMM) protein TIMM17A, a crucial component of the IMM protein import machinery. Excitingly, we also identify the previously uncharacterized effector Lpg2444, as a novel mitochondrially targeted effector, and show that it protects mitochondrial network integrity during mitochondrial stress. Thus, our global proteomics approach is a powerful tool that facilitates the discovery of novel functions and host cell targets of *L.p.* effectors.

## RESULTS

### *L.p.* infection induces T4SS-dependent proteomic changes in the host cell

In order to identify novel host cell components and pathways targeted by *L.p.* effectors, we performed a global proteomics analysis of how *L.p.* infection impacts protein abundance, phosphorylation, and ubiquitination in HEK 293 cells stably expressing the FcγR receptor (HEK293 FcγR cells, **Figure 1A**). This cell line has been used extensively in previous studies to allow efficient internalization of *L.p.* via opsonization with an antibody (Moss et al., 2019; Mukherjee et al., 2011). To discriminate between T4SS-dependent and -independent changes, we utilized wild-type *L.p.* bacteria (WT) or the isogenic *ΔdotA* strain that lacks a functional secretion system (**Figure 1A**). Cells were infected with either of the *L.p.* strains or left uninfected (Ctrl) and lysed after 1h or 8h of infection. The extracted proteins of three biological replicates were trypsinized and separated into 3 aliquots: (1) for the analysis of protein abundance (AB), (2) for the analysis of phosphorylated peptides (phosphoproteome, PH), or (3) for the analysis of ubiquitinated peptides by diglycine (diGly) remnant enrichment which is found upon modification with ubiquitin or the ubiquitin-like proteins NEDD8 and ISG15 (hereafter referred to as ubiquitinome, UB) (Kim et al., 2011; Swaney and Villén, 2016; Udeshi et al., 2012) (**Figure 1A**). Peptides were then subjected to proteomic analysis and quantified (**Table S1** and **Figures S1A-B**). Peptide intensities in biological replicates showed a robust reproducibility with correlation coefficients ranging from 0.79 to 0.93 (**Table S1** and **Figure S1C**). We performed a Principal Component Analysis (PCA) to identify the overall similarities between the different conditions (control, *ΔdotA-1h, ΔdotA-8h,* WT-1h, WT-8h). Interestingly, PCA results captured a larger correlation between control and the mutant strain *ΔdotA* time points relative to the WT time points (**Figure 1B**) for all proteomics assays (especially for PTMs).To compare changes in AB, UB, and PH between the different samples (Ctrl, WT, *ΔdotA),* we calculated the Log_2_ fold changes (Log_2_FC) and the corresponding p-values and adjusted p-values of proteins/proteoforms between WT-infected and uninfected cells (WT/Ctrl), *ΔdotA*-infected and uninfected cells (*ΔdotA*/Ctrl), and WT- and *ΔdotA*-infected cells (WT/*ΔdotA*) using the artMS Bioconductor package (Jimenez-Morales et al., 2019) (**Table S2**). For each pair of relative quantifications, a small fraction of peptides was uniquely identified in only one of the samples while missed in the other one (“missing values”). These peptides might be biologically relevant but could not be represented by the ratio between the two compared samples. To address this issue, artMS enables the relative quantification by assigning (imputing) random intensity values from the limit of MS1 detection to these peptides (Webb-Robertson et al., 2015) and p-values between 0.05 and 0.01 for illustration purposes, but only when identified in one of the conditions in at least 2 or more biological replicates. The imputation allowed us to estimate changes for this interesting category of modified peptides/proteins (see Methods).

**Figure 1:**
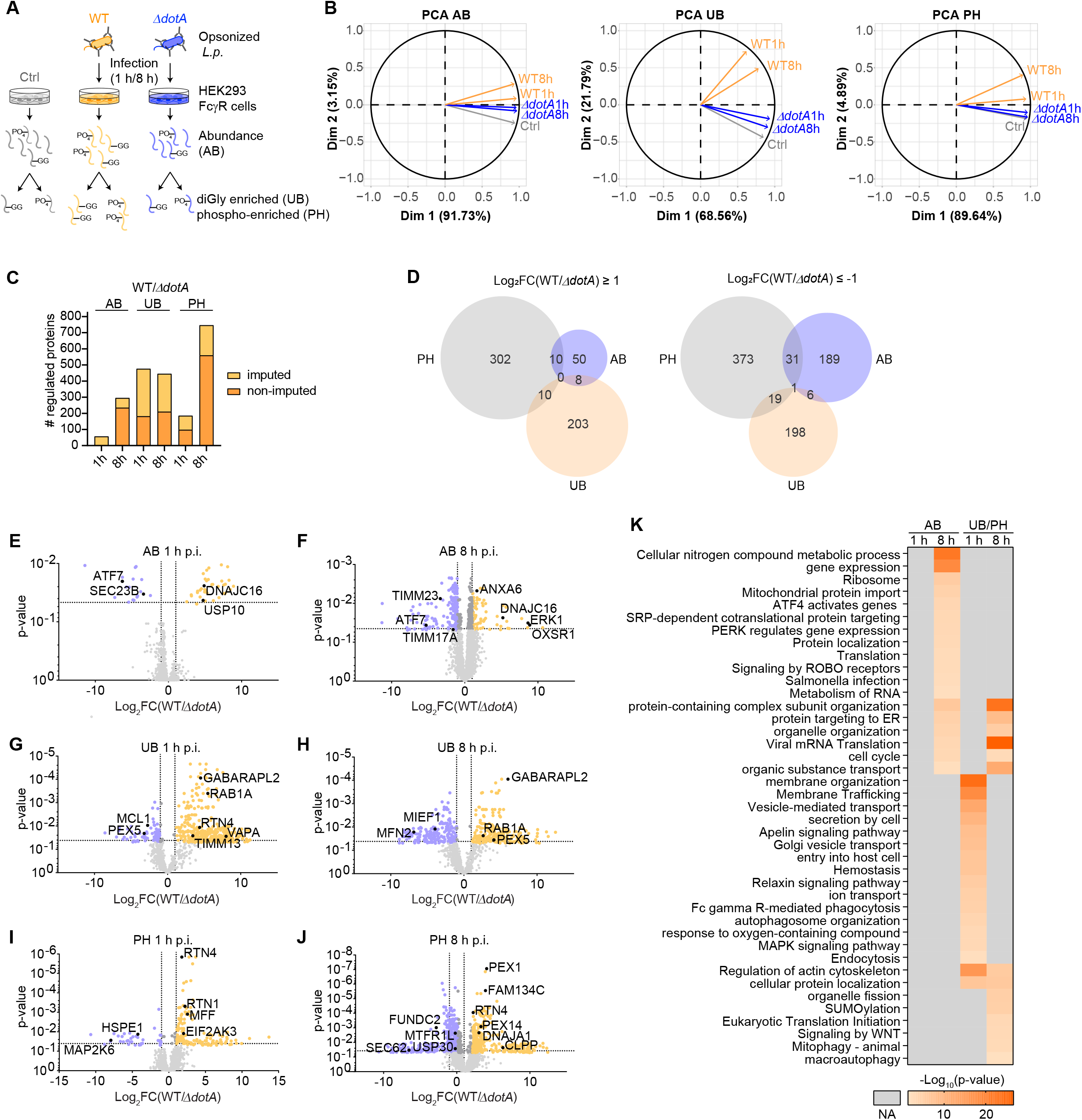
*L.p.* infection induces T4SS-dependent proteomic changes in the host cell. (A) Experimental design. HEK293 FcyR cells were left uninfected (Ctrl) or infected with the opsonized *L.p.* WT or *ΔdotA* strain for 1h or 8h (MOI 100). After tryptic digestion, extracted proteins were directly subjected to MS/MS analysis (abundance, AB) or further enriched for ubiquitinated (UB) and phosphorylated (PH) peptides prior to analysis. (B) Principal Component analysis of normalized MS Intensities of experimental conditions (control, *ΔdotA-1h, ΔdotA-8h,* WT-1h, WT-8h). PC1 and PC2 captured most of the variability. Loading variables are represented as vectors. The smaller angle between control and the mutant time points (*ΔdotA*-1h, *ΔdotA*-8h) implies a larger positive correlation between them, as oppose to a lower correlation (larger angle) between the Control and the WT strain, especially for the later time point (WT-8h). (C) Number of significantly regulated proteins in WT-*vs. ΔdotA*-infected cells (adj.-p-value ≤ 0.05, |Log_2_FC| ≥ 1). Imputed and non-imputed values are highlighted in different shades of orange. (D) Venn diagrams showing the overlap between upregulated (left panel) and downregulated (right panel) proteins in the AB, UB and PH datasets in WT-*vs. ΔdotA*-infected cells (8h p.i.). (E)-(J) Volcano plots showing significantly up-(orange dots) or down-regulated (blue dots) proteins in WT-*vs. ΔdotA*-infected cells (adj.-p-value ≤ 0.05, |Log_2_FC(WT/*ΔdotA*)| ≥ 1). Grey dots: adj.-p-value > 0.05 and/or |Log_2_FC(WT/*ΔdotA*)| < 1. Selected proteins are highlighted in black. (K) Gene ontology enrichment analysis of significantly regulated proteins in WT-*vs*. *ΔdotA*-infected cells (AB, or UB/PH combined) was performed with the g:Profiler g:GO St tool (Raudvere et al., 2019). The heat map shows the most significantly overrepresented GO terms (Biological Processes, Reactome and KEGG pathways).

We found that WT *L.p.* specifically regulated hundreds of proteins as well as hundreds to thousands of modification sites as compared to *ΔdotA* or the uninfected control (cut-off values: adj.-p-value ≤ 0.05, |Log_2_FC| ≥ 1, **Table S3** and **Figures 1C**). The fraction of imputed values was highest in the UB dataset, indicating a possible involvement of bacterial ubiquitin ligases (**Figure 1C**). Importantly, for most proteins, the increase or decrease in protein ubiquitination and phosphorylation was not due to increased or decreased protein abundance as the overlap between regulated proteins in the three datasets was minimal (**Figure 1D**). To understand how the secretion of *L.p.* effectors into the host cell reshapes its proteome, we compared proteins that were significantly up- or down-regulated in WT *vs. ΔdotA*-infected cells (**Figures 1E-J**). As expected, early during infection (1h p.i.), only few proteins were regulated at the level of abundance (**Figures 1C, 1E**, and **Table S3**). However, we observed the WT-specific upregulation of certain proteins such as the co-chaperone DNAJC16 involved in mitochondrial protein import or the deubiquitinase USP10, as well as the down-regulation of few proteins, including the transcription factor ATF7 and SEC23B, a protein involved in ER-to-Golgi transport (**Figure 1E**). The late host proteome response (at 8h p.i.) was characterized by a dramatic decrease in protein abundance in WT-*vs. ΔdotA*-infected cells (**Figure 1F** and **Table S3**). This result was expected as *L.p.* infection is known to inhibit host cell protein synthesis through the combined actions of bacterial effectors (Belyi et al., 2006, 2008; Cornejo et al., 2017; Fontana et al., 2011; Moss et al., 2019; Shen et al., 2009). However, a subset of these down-regulated proteins might also be actively degraded as part of the host cell response to infection or by *L.p.* effectors such as bacterial ubiquitin ligases or proteases. Amongst the down-regulated proteins were TIMM23 and TIMM17, two crucial components of the inner mitochondrial membrane (IMM) protein import machinery. Strikingly, despite the general translation inhibition, the abundance of 68 proteins was increased in WT-infected cells (**Figure 1F** and **Table S3**). These proteins included the oxidative stress-responsive kinase OXSR1, the co-chaperone DNAJC16, the kinase ERK1, and the phospholipid-binding protein Annexin VI (**Figure 1F**). When we compared the ubiquitinome and phosphoproteome in WT *vs. ΔdotA*-infected cells, we found that hundreds of ubiquitinated and phosphorylated proteins were differentially regulated upon WT infection when compared to the *ΔdotA* mutant at both infection time-points (**Figures 1C, 1G-J** and **Table S3**). These proteins included known targets of *L.p.* effectors such as the ER-shaping proteins RTN4 and FAM134C or the small GTPase RAB1 (Horenkamp et al., 2014; Kotewicz et al., 2017; Shin et al., 2020), but also novel hits such as proteins involved in mitochondrial protein import (TIMM13), mitochondrial dynamics and mitophagy (e.g. MIEF1, MFN2, MTFR1L, FUND2C, GABARAPL2), peroxisomal proteins (e.g. PEX1, PEX5, PEX14) and the ER-resident UPR sensor EIF2AK3/PERK (**Figures 1G-J**).

In order to determine which biological processes and pathways were mainly affected upon *L.p.* infection in an T4SS-dependent manner, we performed a functional enrichment analysis of the significantly regulated proteins comparing WT vs. *ΔdotA*-infected cells (AB or UB/PH datasets combined). At the early infection time-point, we did not find any significantly overrepresented terms in the AB dataset (**Figure 1K**). In contrast, at 8h p.i., significantly regulated proteins were enriched for processes and pathways related to proteostasis (e.g. ‘gene expression’, ‘ATF4 activates genes’, ‘PERK regulates gene expression’, ‘protein targeting to the ER’, ‘mitochondrial protein import’) and included additional, unexpected pathways such as ‘signaling by ROBO receptors’. The UB/PH datasets showed significant overrepresentation of expected terms involved in *L.p.* entry into host cells at 1h p.i. (e.g., ‘membrane trafficking’, ‘entry into host cell’, ‘FC gamma R-mediated phagocytosis’, ‘endocytosis’) (**Figure 1K**). At 8h p.i., we found significantly enriched terms involved in organelle homeostasis (e.g., ‘organelle organization’, ‘organelle fission’, ‘mitophagy - animal’). These results reflect the dynamic shift in biological processes and pathways targeted by *L.p.* in a T4SS-dependent manner during the course of infection.

Taken together, *L.p.* induced dynamic changes in host cell protein abundance, ubiquitination, and phosphorylation. Using the *L.p.* WT strain and the *ΔdotA* mutant, we were able to differentiate between a general response to *L.p.* infection and an effector-driven, T4SS-dependent response. Our analysis confirmed previously reported host proteins or pathways targeted by *L.p.* and identified novel proteins/pathways regulated during *L.p.* infection.

### Dynamic regulation of kinase activities during *L.p.* infection

Many cellular stress responses involve the activation of kinases through phosphorylation events. We used the open source tool PhosFate Profiler (Ochoa et al., 2016) to infer host cell kinase and complex regulation in WT-*vs. ΔdotA*-infected cells based on the identified, significantly regulated phosphorylation sites. Visualization of the inferred kinase activities on the kinome tree with CORAL (Metz et al., 2018) revealed the regulation of all main kinase groups in response to *L.p.* WT infection, with the exception of the TK kinase family in which only few kinases were regulated (**Figures 2A, S2A** and **Table S4**). About one third of the kinases were differentially regulated at 1h p.i. and 8h p.i., highlighting the dynamic nature of the kinase-mediated host cell response to the pathogen. For instance, cyclin-dependent kinase 2 (CDK2) showed increased activity at 1h p.i., but decreased activity at 8h p.i. (**Figures 2A-B** and **S2A**). A similar trend was observed for BRSK2, a kinase involved in the regulation of cell cycle and ER-stress mediated apoptosis (Wang et al., 2012). At both infection time-points, the kinase triade VRK1-PLK3-ERK7, which plays a role in Golgi disassembly (López-Sánchez et al., 2009), showed higher activity in WT-*vs. ΔdotA*-infected cells (**Figures 2A-B** and **S2A**). Similarly, the activity of the Jun kinase JNK2 was elevated at 1h p.i. and 8h p.i.. Accordingly, we observed the phosphorylation of its target c-Jun at serine 73 (Jaeschke et al., 2006; Kallunki et al., 1994) during the course of WT-infection, but not in *ΔdotA*-infected cells (**Figure 2C**). Functional enrichment analysis of all kinases with predicted increased activity in WT *L.p*.-infected cells at both infection time-points revealed the significant overrepresentation of selected pathways/processes such as ‘cellular response to stress’, ‘apoptotic process’, ‘positive regulation of protein metabolic process’, ‘cytoskeleton organization’, ‘mitophagy-animal’ and ‘sphingolipid signaling pathway’ (**Figure 2D**).

**Figure 2:**
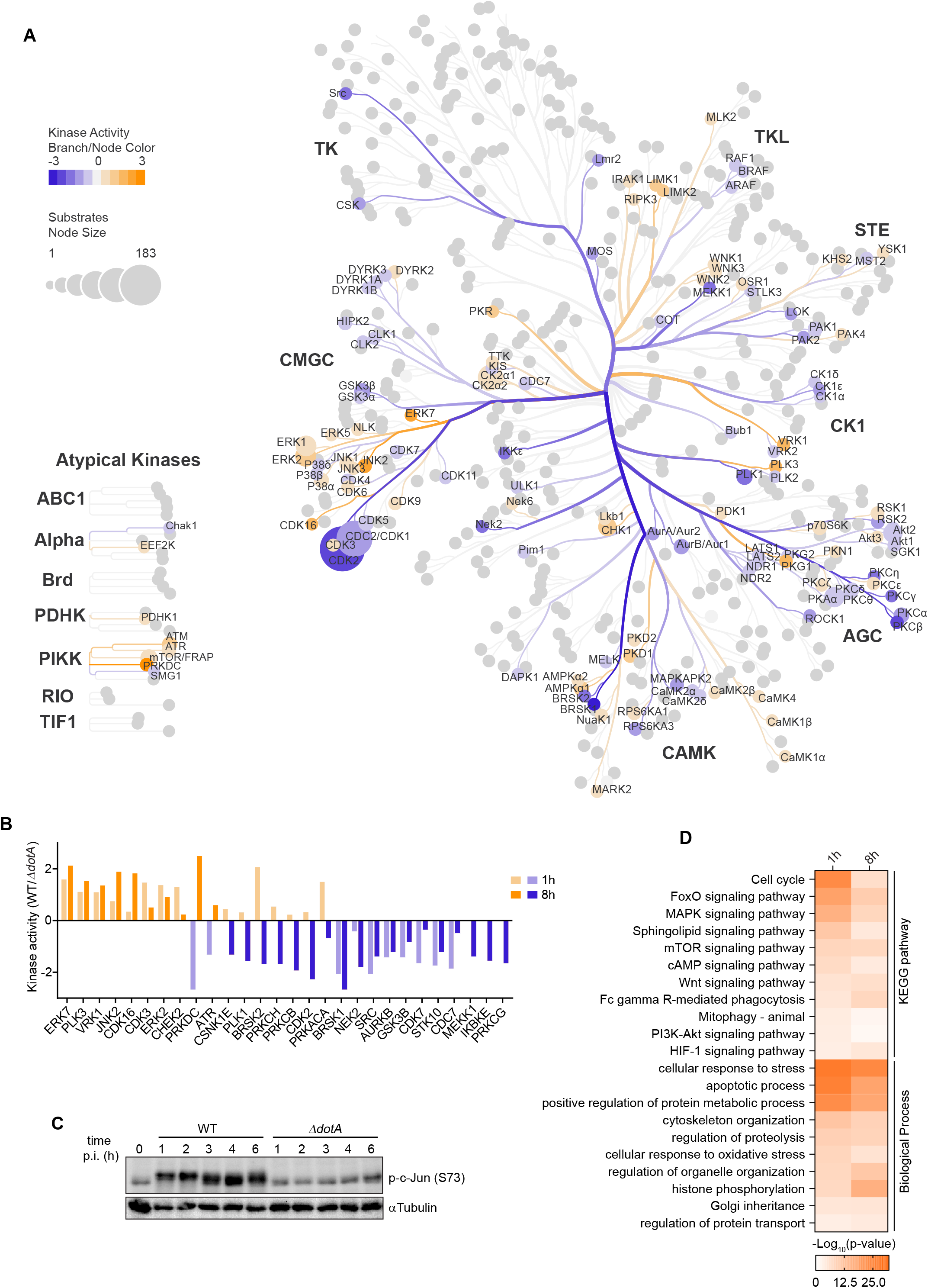
Dynamic regulation of kinase activities during *L.p.* infection. (A) Host cell kinase activities in WT-*vs*. *ΔdotA*-infected cells at 8h p.i. were inferred with PhosFate Profiler (Ochoa et al., 2016) based on regulated phosphorylated sites, and mapped on the kinase tree with CORAL (Metz et al., 2018). Kinase activity is indicated by the branch and node color, and the number of substrates by the node size. Names of kinase families and regulated kinases are highlighted. (B) Inferred kinase activities of the most significantly regulated kinases (p-value ≤ 0.01) in WT-*vs*. *ΔdotA*-infected cells at 1h p.i. and 8h p.i.. (C) Immunoblot analysis of phosphorylated c-Jun at serine 73 (p-c-Jun (S73)) and αTubulin (loading control) levels in HEK293 FcγR cells during the course of infection with the *L.p.* WT-or *ΔdotA* strain (MOI 100). (D) Gene ontology enrichment analysis of kinases with predicted upregulated activities in WT-*vs*. *ΔdotA*-infected cells. Shown are the most significantly overrepresented pathways (KEGG pathways) and biological processes.

The analysis of the phosphoproteome with PhosFate Profiler also allows the prediction of the activities of co-regulated protein complexes (Ochoa et al., 2016). Similar to the inferred kinase activities, some protein complexes showed a comparable regulation early and late during infection, including the stress-induced, transcriptional ERα-c-Jun complex (**Figure S2B** and **Table S4**). However, most protein complexes were regulated in an opposite manner at 1h p.i. and 8h p.i.. For instance, the RNF20-RNF40-UbE2E1 complex which mediates histone monoubiquitination (Zhu et al., 2005), shows decreased activity in WT-infected cells at 1h p.i., but increased activity at 8h p.i. (**Figure S2B**). In contrast, the BMI1-HPH1-HPH2 complex involved in repression of gene expression (Alkema et al., 1997) was predicted to be highly active at 1h p.i., but down-modulated at 8h p.i.. These results give new insights into the dynamics of host cell kinase signaling and complex regulation in response to *L.p.* infection and highlight new biological pathways and processes that might be exploited by *L.p.* effectors.

### Spatio-temporal changes of the host cell proteome, ubiquitinome and phosphoproteome in response to *L.p.* WT infection

In order to gain a spatio-temporal overview of the T4SS-dependent proteomics changes during *L.p.* infection, we mapped the numbers of regulated proteins (AB, UB, and PH combined) on their predicted major subcellular localization according to their primary evidence code (ECO), or, if not available, to their documented subcellular localization in The Human Protein Atlas database (http://www.proteinatlas.org) (Thul et al., 2017). This analysis revealed that the predicted subcellular localizations with the highest levels of protein regulation were the nucleus, the cytosol and the cell membrane both early and late during *L.p.* infection (100-500 regulated proteins, **Figures 3A-B).** With the exceptions of endosomes and the Golgi apparatus, the number of regulated proteins increased in all subcellular localizations during the course of infection. The most dramatic increase was induced in the nucleus (307 proteins), the cytoplasm (142 proteins), the nucleolus (48 proteins) and in mitochondria (35 proteins) (**Figures 3A-B**). The nucleolus has recently been implicated in the resistance to bacterial pathogens and therefore resembles an attractive target for intracellular bacteria (Tiku et al., 2018). Interestingly, the *L.p.* effector RomA/LegAS4 has been proposed to modulate nucleolar function through the modification of rDNA chromatin (Li et al., 2013; Rolando et al., 2013), further suggesting that the nucleolus is rewired during *L.p.* infection through the actions of one or more bacterial effectors and/or as part of the host cell response. Our study thus might serve as a platform to gain more insights into the regulation of these newly emerging organelles targeted by *L.p.* effectors.

**Figure 3:**
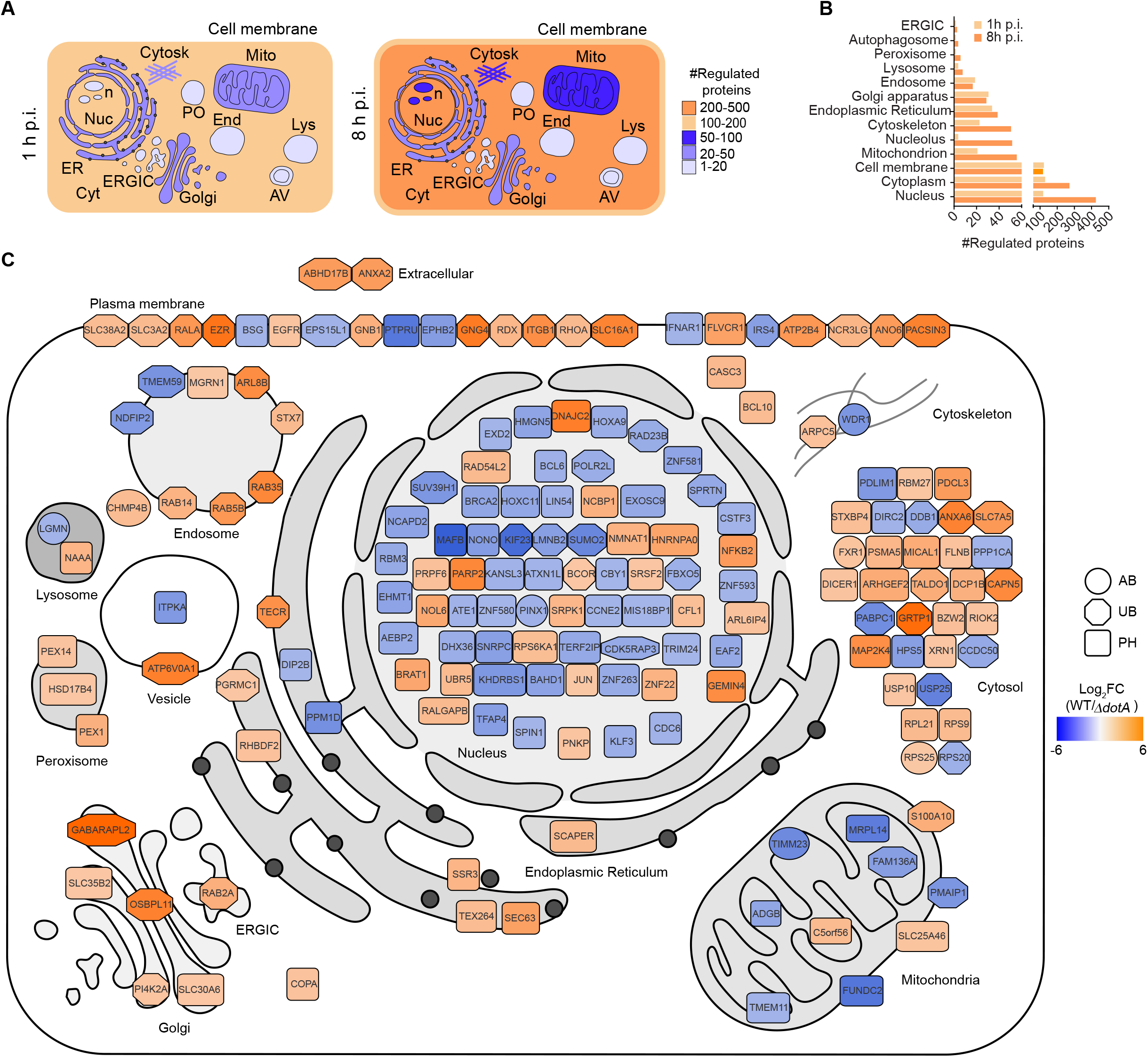
Spatio-temporal changes of the host cell proteome, ubiquitinome and phosphoproteome in response to *L.p.* WT infection. (A) The numbers of regulated proteins in WT *vs. ΔdotA* infected cells (AB, UB and PH combined) were quantified for and mapped on each predicted subcellular compartment. The number range of regulated proteins is indicated by the color code. Cytosk: cytoskeleton, Mito: mitochondria, PO: peroxisomes, n: nucleolus, Nuc: nucleus, ER: endoplasmic reticulum, ERGIC: ER-Golgi intermediate compartment, Cyt: cytosol, AV: autophagosome, End: endosome, Lys: lysosome. (B) Bar graph showing quantifications of panel (A). (C) Snapshot of WT-infected calls at 8h p.i.. Highly significantly regulated proteins (adj.-p-value ≤ 0.01, |Log_2_FC(WT/*ΔdotA*)| ≥ 2) were mapped on the predicted host cell organelles according to their primary ECO with Cytoscape (Shannon et al., 2003). The Log_2_FC values are indicated by a color scale (orange: up-regulated, blue: down-regulated), the dataset is indicated by the shape of the icon (circle: AB, octagon: UB, rounded square: PH).

To obtain a more detailed picture of how host cell organelles are rewired at each infection time-point, we mapped the top regulated proteins at 1h p.i. (adj.-p-value ≤ 0.001, (|Log_2_FC(WT/*ΔdotA*)| ≥ 1) and 8h p.i. (adj.-p-value ≤ 0.001, (|Log_2_FC(WT/*ΔdotA*)| ≥ 2) on the predicted host cell organelles (**Figures S3** and **3C**). After *L.p.* is phagocytosed by the host cell, the secretion of bacterial effectors prevents fusion with the endo-lysosomal compartment and leads to the recruitment of ER-derived vesicles that are required for the establishment of the LCV (Cornejo et al., 2017; Qiu and Luo, 2017b). In agreement with *L.p.’s* progression through the host cell, the top hits at 1h p.i. were mainly located in the cell membrane, the endosome, the ER and the Golgi apparatus (**Figure S3**). Most of these top-regulated proteins (125/138) showed increased ubiquitination or phosphorylation in response to *L.p.* WT infection. Some of the hits were expected as they are known to be modified by *L.p.* effectors, such as the GTPases RAB1A, RAB35 and components of the ARP2/3 complex (Michard and Doublet, 2015; Mukherjee et al., 2011; Qiu et al., 2016). Other proteins, such as the vesicle-trafficking protein SEC22B, have been shown to be required for LCV formation (Arasaki and Roy, 2010; Kagan et al., 2004), but a modification of these proteins in response to infection has not been documented. In addition, the ER-shaping ER-shaping/ER-phagy proteins RTN1, RTN4, ATL3 and FAM134C were among the top regulated proteins and showed increased phosphorylation upon *L.p.* infection. These proteins were recently identified as phosphoribosyl-ubiquitinated targets of the non-canonical *L.p.* ubiquitin ligase SdeA (Shin et al., 2020), suggesting a possible crosstalk with protein phosphorylation. Interestingly, we found modified proteins that have not been linked to *L.p.* infection but might play a role for bacterial replication and/or be targets of *L.p.* effectors. For instance, proteins involved in mitochondrial membrane organization and ER-mitochondria tethering were highly regulated upon *L.p.* infection (MFF, PMAIP1, SLC25A46, VAPB, MMGT1). In contrast to the early infection time-point, the top hits at 8h p.i.- a time-point where the LCV is established and resembles the rough ER-included less cell membrane proteins and showed a broader distribution throughout the cell (**Figure 3C**). Most of the top regulated proteins were predicted to be in the nucleus (67/196). In accordance with the important role of the ER for *L.p.* replication, several ER proteins were amongst the highly regulated proteins. Interestingly, the recently discovered ER-phagy receptor TEX264, which is also an SdeA mediated phosphoribosyl-ubiquitin target, (An et al., 2019; Chino et al., 2019; Shin et al., 2020) was modified by phosphorylation in WT-infected cells, further highlighting a potential role of ER-phagy during *L.p.* infection. Also, the ER translocon component SEC63 showed increased phosphorylation which correlates with the recruitment of the rough ER at this time-point and might hint towards an unprecedented function of this protein for *L.p.* replication. Analogous to the early infection time-point, several mitochondrial proteins were among the top hits at 8h p.i. (TIMM23, FAM136A, ADGB, MRPL14, C5orf56, TMEM11, PMAIP1, S100A10, FUNDC2, SLC25A46). Taken together, the spatio-temporal analysis of host cell proteome changes during *L.p.* infection provides novel insights into the dynamic targeting of host cell organelles by *L.p..*

### *L.p.* reshapes the mitochondrial proteome

Our proteomics analysis highlighted mitochondria as one of the targets of *L.p.* infection. A few reports discuss host cell mitochondria as a potential target of *L.p.* (Arasaki et al., 2017; Banga et al., 2007; Degtyar et al., 2009; Dolezal et al., 2012; Escoll et al., 2017; Laguna et al., 2006; Tiku et al., 2020), but the exact functional role of mitochondria during infection and possible modifications of mitochondrial proteins by *L.p.* effectors remain largely elusive. We therefore set out to analyze the effects of *L.p.* on the regulation of proteins with predicted mitochondrial localization in more detail. When we compared the mitochondrial proteome in WT- and *ΔdotA*-infected cells, we found that mitochondrial proteins were significantly regulated in all three datasets and that the number of regulated proteins increased during the course of infection (**Figure 4A**). Functional enrichment analysis of all regulated mitochondrial proteins in WT-*vs*. *ΔdotA*-infected cells showed that at 1h p.i., proteins involved in mitochondrial membrane organization, regulation of cell death, mitochondrial protein import, mitochondrial fission and fusion or with chaperone functions were significantly enriched (**Figures S4A-B**). This early response to WT *L.p.* infection included predominantly ubiquitination and phosphorylation events such as the increased ubiquitination of the mitochondrial chaperone HSPD1 or phosphorylation of the mitochondrial fission factor MFF (**Figure S4B**). At 8h p.i., the most significantly overrepresented terms were mitochondrial protein import, mitochondrial gene expression, oxidative phosphorylation, mitochondrial fission and fusion, response to unfolded protein, apoptotic signaling pathway, Clp/crotonase-like domain superfamily and chaperones (**Figures 4B-C**). Several components of the mitochondrial protein import (e.g. TIMM17A, TIMM23) and gene expression machineries (e.g. MRPS7, MRPL14) were down-regulated at the level of protein abundance in WT-infected cells (**Figure 4C**). Additionally, multiple proteins were regulated by PTMs. For instance, the translocases of the outer mitochondrial membrane (OMM), TOMM22 and TOMM70, were differentially phosphorylated at multiple sites which is thought to regulate mitochondrial protein import (Matic et al., 2018; Schmidt et al., 2011). The down-modulation of mitochondrial protein import and mitochondrial gene expression are hallmarks of mitochondrial stress responses such as the UPR^mt^ (D’Amico et al., 2017). Thus, we show for the first time that *L.p.* infection reshapes the mitochondrial proteome in a T4SS-dependent manner, mainly affecting proteins involved in mitochondrial proteostasis and mitochondrial dynamics.

**Figure 4:**
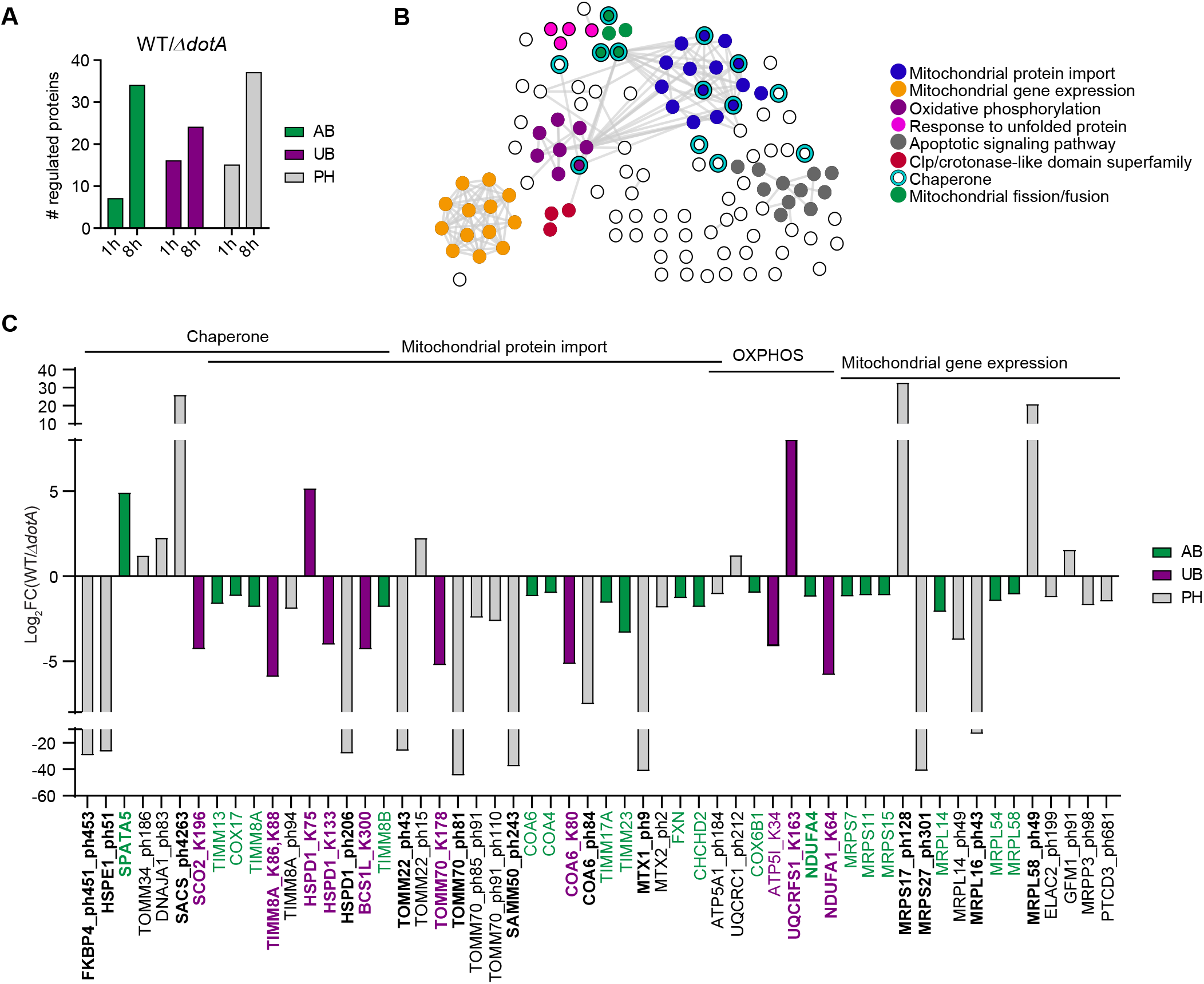
*L.p.* reshapes the mitochondrial proteome. (A) Percentage of mitochondrial proteins/proteoforms whose abundance (AB), ubiquitination (UB) or phosphorylation (PH) was regulated in WT-*vs. ΔdotA*-infected cells, (adj.-p-value ≤ 0.05, |Log_2_FC| ≥ 1) at 1h p.i. and 8h p.i.. (B) Gene ontology enrichment and network analysis of all regulated mitochondrial proteins in WT-*vs*. *ΔdotA*-infected cells at 8h p.i. (AB, UB, PH combined) was performed with the Cytoscape stringApp (Doncheva et al., 2019; Shannon et al., 2003). Each circle represents one protein. Selected overrepresented pathways are highlighted and annotated. (C) Log_2_FC(WT/*ΔdotA)* values of significantly regulated mitochondrial proteins/proteoforms from selected gene ontology terms in (E). Shown are changes in protein abundance (green), ubiquitination sites (purple) and phosphorylation sites (grey) at 8h p.i.. Proteins/proteoforms with imputed values are highlighted in bold.

### *L.p.* infection induces a mitochondrial stress response

Earlier work has shown that infection of the nematode *Caenorhabditis elegans (C.elegans)* with pathogenic *Pseudomonas aeruginosa* induces the expression of UPR^mt^ genes and that chemical activation of the UPR^mt^ results in the induction of innate immunity genes, thereby providing resistance of the animal to the pathogen (Pellegrino et al., 2014). However, the role of the UPR^mt^ during bacterial infection in mammalian cells is not known. Given our observation that proteins involved in mitochondrial proteostasis are regulated upon *L.p.* infection (**Figures 4** and **S4**), we set out to investigate if *L.p.* induces and/or modulates the UPR^mt^. Comparison of significantly regulated proteins/proteoforms in response to *L.p.* WT or *ΔdotA* infection at 8h p.i. with a published list of UPR^mt^-regulated genes (Münch and Harper, 2016) (adj.-p-value ≤ 0.05, |Log_2_FC(Treatment/Ctrl)| ≥ 0.6, threshold based on (Münch and Harper, 2016)) revealed that a substantial number of UPR^mt^-related proteins/proteoforms were regulated upon *L.p.* infection (**Figure 5A**). Except for ubiquitinated proteins, the number of overlapping proteins was 6-10 times higher in WT-infected cells when compared to the *ΔdotA* mutant, indicating a strong contribution of bacterial effectors to the regulation of UPR^mt^ proteins. Amongst the UPR^mt^ proteins that were up-regulated during *L.p.* WT infection were the two transcription factors ATF3 and JUN, proteins involved in RNA transport or splicing (FXR1, ARL6IP4, CWF19L2), and GFER, a sulfhydryl oxidase important for oxidative folding of proteins in the mitochondrial intermembrane space (IMS) (**Figure 5B**, orange bars). The UPR^mt^ proteins that were down-regulated in *L.p.* infection included several RNA-binding or -processing proteins (DIEXF, SKIV2L2, ZNF638, CSTF3), protein phosphatase subunits (PPP1R10, PPP1CC) and proteins involved in mitochondrial protein import (TIMM17A, TOMM5) (**Figure 5B**, purple bars). The T4SS-dependent induction of ATF3 and the down-regulation of TIMM17A during *L.p.* infection was validated by immunoblot and correlated with the processing of the long isoform of OPA1 (L-OPA-1) (**Figure 5C**) and the increased phosphorylation of c-Jun at S73 (**Figure 2C**), indicative of the activation of a mitochondrial stress response (Ahola et al., 2019; Jovaisaite et al., 2014).

**Figure 5:**
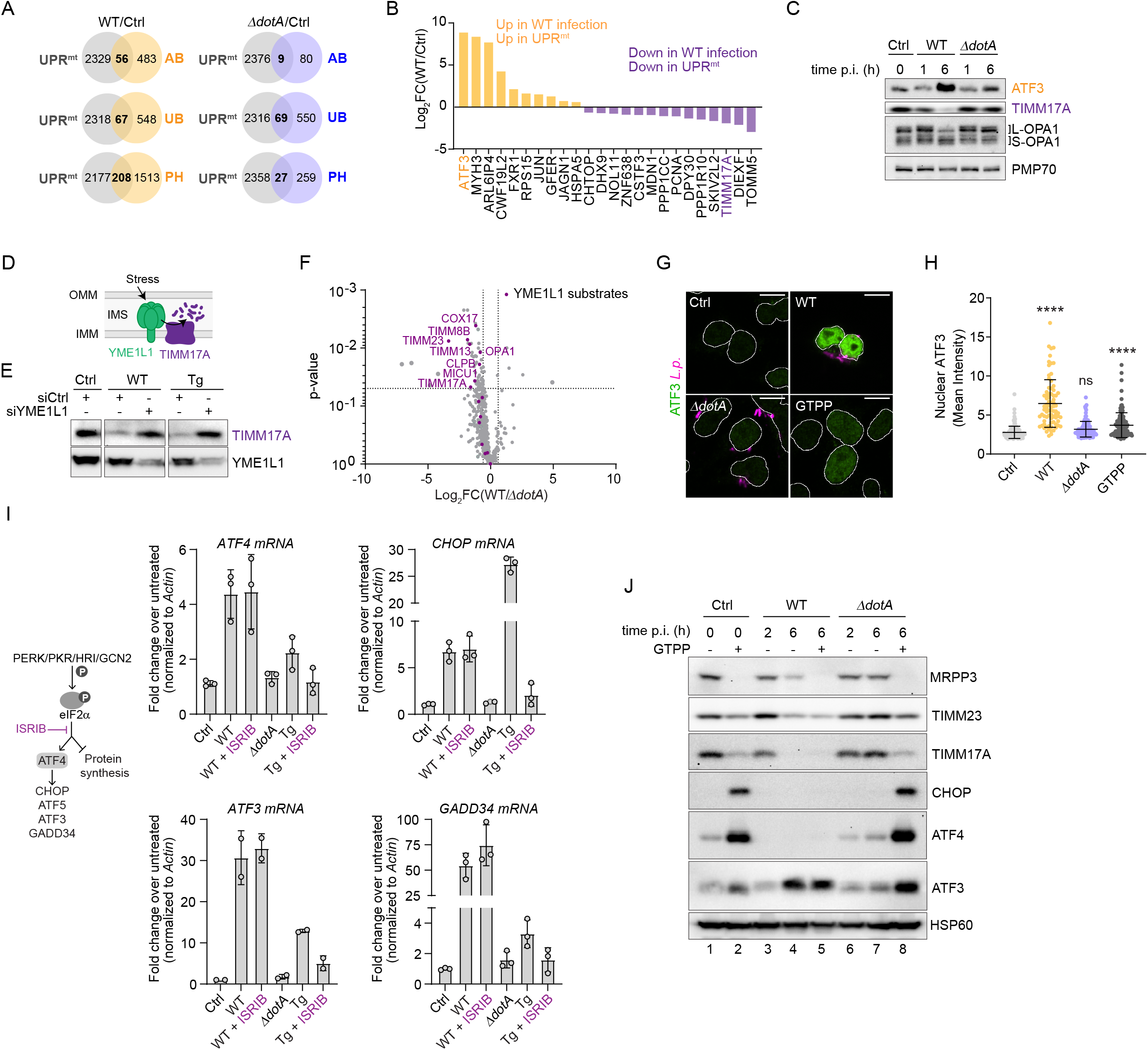
*L.p.* infection modulates the mitochondrial stress response. (A) Comparison of significantly regulated proteins/proteoforms in response to *L.p.* WT or *ΔdotA* infection (at 8h p.i.) with a published list of UPR^mt^-regulated genes (Münch and Harper, 2016). The Log_2_FC threshold was adjusted according to the published dataset (adj.-p-value ≤ 0.05, |Log_2_FC| ≥ 0.6). Shown is the overlap between the different datasets in WT-infected *vs.* Ctrl cells (left panel) or *ΔdotA*-infected *vs.* Ctrl cells (right panel). (B) Bar graph showing Log_2_FC(WT/Ctrl) values in abundance of selected proteins that are either up-or down-regulated in both UPR^mt^ and WT-infected cells (adj.-p-value ≤ 0.05). ATF3 and TIMM17A are highlighted. (C) HEK293 FcγR cells were left uninfected (Ctrl) or infected with *L.p.* WT or *L.p. ΔdotA* for 1h and 6h (MOI 25). Protein levels of ATF3, TIMM17A, OPA1 (control for mitochondrial stress) and PMP70 (loading control) were analyzed by immunoblot. L-OPA1: long OPA-1, S-OPA1: short OPA-1. n=2 biological replicates. (D) During different stress conditions, the i-AAA protease YME1L1 located in the IMM is activated and cleaves TIMM7A to decrease mitochondrial protein import. (E) HEK293 FcyR cells were transfected with control siRNA or siRNA targeting YME1L1 and either left untreated, infected with WT *L.p.* (MOI 25) or treated with 10 μM thapsigargin (Tg, positive control) for 6h. TIMM17A and YME1L1 protein levels were analyzed by immunoblot. (F) Volcano plot showing all detected mitochondrial proteins WT-*vs. ΔdotA*-infected cells. Known stress-induced YME1L1 substrates (MacVicar et al., 2019) are highlighted in purple. (G) The levels of nuclear ATF3 in untreated (Ctrl), GTPP-treated (10 μM, 6h), *L.p.* WT- or *Δdot*-infected HEK293 FcyR cells (MOI 5, 6h) were analyzed by immunofluorescence microscopy. Cells were stained with Hoechst33342 to define the nuclear area (shown as outline), anti-ATF3 antibody (green) and anti-*L.p.* antibody (magenta). Scale bar: 10 μm. (H) The background corrected, nuclear ATF3 signal (mean intensity) was quantified by automated image analysis with CellProfiler (Carpenter et al., 2006). Each dot represents one cell. Shown is the mean ± SD of n ≥ 78 cells. Statistical differences were analyzed by one-way ANOVA and Tukey’s multiple comparison test. p-value **** p ≤ 0.0001, p-value ** p ≤ 0.01, ns p > 0.05. (I) The cartoon on the right depicts signaling through the ISR (described in the text). HEK293 FcyR cells were infected with L.p. WT or *ΔdotA* (6h infection, MOI 100) in presence or absence of the inhibitor ISRIB (200 nM) for 6h. Thapsigargin (Tg, 10 μM) was used as a positive control. qPCR of *ATF4, CHOP, ATF3* and *GADD34* mRNA levels were analyzed by qPCR. Shown are the mean levels relative to the control ± SD of n = 3 or n = 2 (*ATF3*) biological replicates. (J) Uninfected (Ctrl), WT- or *ΔdotA*-infected (MOI 25) HEK293 FcyR cells were treated with DMSO (-) for 2h or 6h, or GTPP (+, 10 μM) for 6h. Protein levels of the UPR^mt^ markers MRPP3, TIMM23, TIMM17A, CHOP, ATF4, and ATF3 (loading control: HSP60) were analyzed by immunoblot.

Both OPA-1 and TIMM17A are known substrates of the mitochondrial i-AAA protease YME1L1 which is activated during certain stress conditions such as mitochondrial or nutrient stress, thereby reshaping the mitochondrial proteome (MacVicar et al., 2019; Rainbolt et al., 2013) (**Figure 5D**). Therefore, we tested whether the down-regulation of TIMM17A during *L.p.* WT infection is due to general, effector-mediated inhibition of host cell translation (Belyi et al., 2006, 2008; Cornejo et al., 2017; Moss et al., 2019; Shen et al., 2009), or is actively mediated by YME1L1. In both *L.p.* infected cells and cells treated with thapsigargin (Tg), a known inducer of YME1L1 (Rainbolt et al., 2013), TIMM17A levels were decreased (**Figure 5E**). However, siRNA-mediated knockdown of YME1L1 stabilized TIMM17A under both conditions to a similar extent, suggesting that the downregulation of TIMM17A during *L.p.* WT infection requires active YME1L1. Furthermore, the protein abundance of several other recently discovered, stress-induced YME1L1 substrates (MacVicar et al., 2019) was down-regulated in *L.p.* WT infected cells (**Figure 5F**), suggesting a potential novel role of YME1L1 in the response to *L.p.* infection.

ATF3 is a bZIP domain transcription factor implicated in multiple cellular stress responses that can either activate or repress transcription of its target genes (Jadhav and Zhang, 2017). Interestingly, ATF3 has been recently discussed as one of the putative mammalian homologues of ATFS-1, the central UPR^mt^ regulator in *C.elegans* (Münch and Harper, 2016). ATF3 was also shown to repress the transcription of PINK1, an important player in mitophagy, further highlighting its potential role for mitochondrial homeostasis (Bueno et al., 2018). Consistently, nuclear ATF3 was significantly enhanced in cells treated with gamitrinib-triphenylphosphonium (GTPP), a specific inhibitor of the mitochondrial chaperone HSP90/TRAP1 and inducer of the UPR^mt^ (Kang et al., 2009; Münch and Harper, 2016; Siegelin et al., 2011) (**Figures 5G-H**). Nuclear ATF3 induction was even higher in cells infected with the *L.p.* WT strain, but not significantly induced in *ΔdotA*-infected cells. In addition, *ATF3* mRNA was dramatically induced upon *L.p.* infection in a T4SS-dependent manner (**Figure S5A**). Similarly, the transcripts of other putative mammalian ATFS-1 homologs, *ATF4, CHOP* and *ATF5* (Fiorese et al., 2016; Quirós et al., 2017) as well as their down-stream target *GADD34* were elevated in response to *L.p.* WT, but not *ΔdotA* infection (**Figure S5A**). Furthermore, the transcript induction of *ATF4* and *CHOP* increased with an increasing multiplicity of infection (MOI) in WT-infected cells (**Figure S5B**). These data suggested that *L.p.* WT infection activates the ISR. Activation of the ISR kinases (PERK, PKR, GCN and HRI) results in the phosphorylation of eIF2α at serine 51, thereby causing a global translational attenuation while allowing the preferential translation of the transcription factor ATF4 and its downstream targets (Pakos-Zebrucka et al., 2016) (cartoon in **Figure 5I**). To test whether ISR activation was required for stress gene induction during *L.p.* infection, we treated the cells with or without the ISR inhibitor ISRIB (Sidrauski et al., 2013) (cartoon in **Figure 5I**) and analyzed the mRNA levels of *ATF3, ATF4, CHOP* and *GADD34.* Much to our surprise, ISRIB did not prevent the induction of these transcripts upon infection with *L.p.* **(Figure 5I**). In contrast, the induction of the same genes upon treatment with Tg was almost completely abolished by ISRIB. Thus, signaling through the ISR is not required for the transcriptional induction of *ATF3, ATF4, CHOP* and *GADD34* during *L.p.* infection. In order to get a more detailed understanding of the regulation of the mitochondrial stress response during *L.p.* infection, we compared the protein levels of several UPR^mt^ related proteins in uninfected, WT- or *ΔdotA*-infected cells in absence or presence of GTPP by immunoblot analysis. As expected, the induction of the UPR^mt^ with GTPP led to the degradation of the mitochondrial ribonuclease P catalytic subunit MRPP3 (Münch and Harper, 2016) (**Figure 5J**, lane 2 *vs.* 1). Similarly, infection with the *L.p.* WT strain, but not the *ΔdotA* strain, led to a robust reduction in MRPP3 levels after 6h p.i. (**Figure 5J**, lane 4 *vs.*1), which was further reduced upon additional treatment with GTPP (**Figure 5J**, lane 5 *vs.*1). Likewise, TIMM17A and TIMM23 protein levels were reduced in both GTPP-treated and WT-infected, but not in *ΔdotA-* infected cells. Furthermore, GTPP led to a strong increase in ATF4, CHOP and ATF3 protein levels in uninfected and *ΔdotA*-infected cells (**Figure 5J**, lane 2 *vs.* 1 and lane 8 *vs.* 6). Strikingly, the GTPP-mediated induction of CHOP and ATF4 at the protein level was completely suppressed in WT-infected cells (**Figure 5J**, lane 5 *vs*. 2). In contrast, ATF3 was strongly induced in untreated and GTPP-treated cells infected with the WT strain, suggesting an effector-mediated, selective modulation of this stress response.

In short, *L.p.* induces an effector-driven mitochondrial stress response that shares similarities with the UPR^mt^ and is characterized by: i) the down-modulation of the IMM protein import machinery, at least partially regulated by the protease YME1L1, ii) the ISR-independent up-regulation of stress-related transcripts and iii) the selective induction of ATF3 at the protein level. However, in contrast to the UPR^mt^, *L.p.* infection suppresses the translation of ATF4 and CHOP, two central transcription factors of ISR-dependent stress responses, thereby modulating the translational output of the UPR^mt^.

### The *L.p.* effector Lpg2444 is targeted to host cell mitochondria and has mitoprotective functions

Most of the changes in the mitochondrial proteome upon *L.p.* infection were dependent on a functional T4SS (**Figure 4**), suggesting the direct or indirect involvement of *L.p.* effector proteins. Few reports have shown that certain *L.p.* effectors can translocate into the host cell mitochondria (Degtyar et al., 2009; Dolezal et al., 2012). To predict whether additional *L.p.* effectors might target host cell mitochondria, we performed a computational analysis of all known *L.p.* effectors using the complementary tools MitoFates, MitoProt, TargetP1.1, DeepLoc and CELLO (Almagro Armenteros et al., 2017; Claros and Vincens, 1996; Emanuelsson et al., 2000; Fukasawa et al., 2015; Sun and Habermann, 2017; Yu et al., 2006). Our analysis identified a total of 18 *L.p.* effector proteins with a high probability of mitochondrial localization (**Figure 6A**). As a proof of concept, two of these identified effectors, Lpg2905 and Lpg2176/LegS2, have been previously reported to localize to host cell mitochondria (Degtyar et al., 2009; Dolezal et al., 2012). We generated GFP-tagged versions of a subset of effectors to confirm their mitochondrial localization upon ectopic expression in HeLa FcγR and HEK293 FcyR cells. As expected, Lpg2176/LegS2 co-localized with the mitochondrial marker TOMM20 (**Figure S6A**). Interestingly, Lpg2444, a previously uncharacterized effector, also showed significant mitochondrial localization (**Figures 6B-D** and **S6A**). Subcellular fractionation of cells transiently transfected with GPF or GFP-Lpg2444 confirmed that the effector almost exclusively localized to host cell mitochondria, while GFP was mainly found in the cytosolic fractions (**Figure 6E**). Thus, our data show that the computational prediction of mitochondrial localization of *L.p.* effectors is a promising approach to identify novel bacterial effectors that can localize to host cell mitochondria.

**Figure 6:**
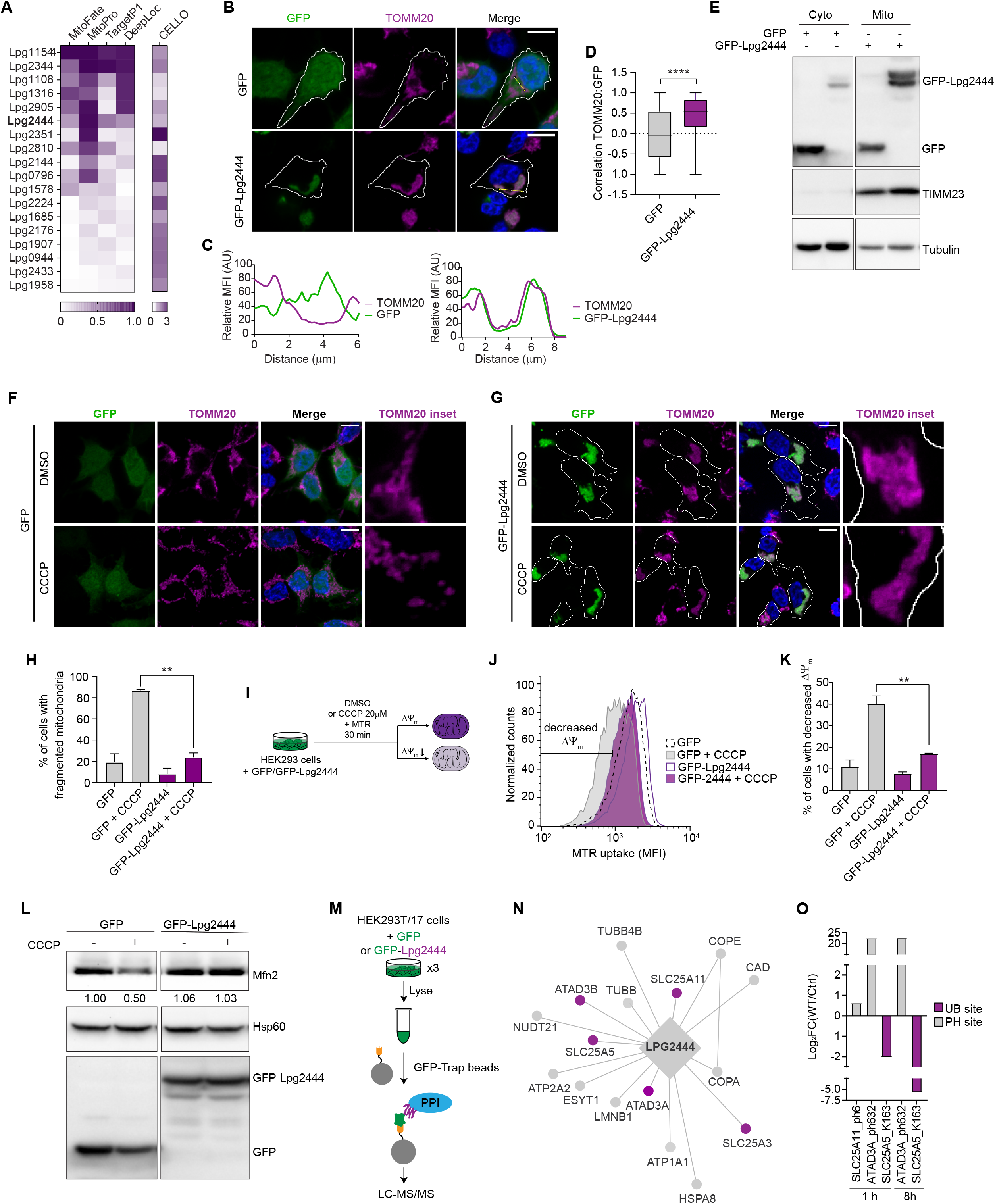
The *L.p.* effector Lpg2444 is targeted to host cell mitochondria and protects mitochondrial network integrity. (A) Computational analysis of all known *L.p.* effectors showed a high probability of mitochondrial localization for 18 *L.p.* effectors. Shown are the scores obtained with the different prediction algorithms as indicated. The uncharacterized effector Lpg2444 is highlighted in bold. (B) HEK293 FcyR cells were transfected with GFP or GFP-Lpg2444 and analyzed by immunofluorescence microscopy. Mitochondria were stained with an antibody against the OMM protein TOMM20 (magenta) and nuclei were stained with Hoechst33342 (blue). Scale bar: 10 μm. (C) Line profiles as depicted in the merged images in (B) showing the correlation of normalized GFP (green) and TOMM20 (purple) median fluorescence intensities (MFI) in GFP-transfected (left panel) or GFP-Lpg2444-transfected cells (right panel). (D) The correlation (Pearson’s coefficient) between the green (GFP) and far red (TOMM20) channel was quantified in an automated fashion with CellProfiler (Carpenter et al., 2006). Shown is a box plot of the values derived from n > 900 transfected cells. Two-tailed p-value **** p ≤ 0.0001. (E) HEK293 FcyR cells transiently transfected with GFP or GFP-Lpg2444 were subjected to subcellular fractionation. Cytosolic and mitochondrial fractions were analyzed by immunoblot using antibodies against GFP, TIMM23 (mitochondrial control) and Tubulin (cytosolic control). (F-H) Immunofluorescence analysis of mitochondrial morphology in GFP-transfected (F) or GFP-Lpg2444-transfected HEK293 FcyR cells (G) treated with DMSO or 10 μM CCCP for 6h. Mitochondria were stained with an anti-TOM20 antibody (magenta) and nuclei were stained with Hoechst33342 (blue). Scale bar: 10 μm. Cells were scored according to their mitochondrial morphology (fragmented *vs.* non-fragmented) and the percentage of cells with fragmented mitochondria was quantified from a total number of n ≥ 100 cells (H). Shown are the mean values ± SD of n = 2 technical replicates. Statistical differences were analyzed by one-way ANOVA and Tukey’s multiple comparison test. ** p ≤ 0.01. (I-K) Analysis of the effect of Lpg2444 on ΔΨm. GFP- or GFP-Lpg2444 transfected HEK293 FcγR cells were treated with DMSO or 20 μM CCCP in addition to 200 nM Mitotracker Red FM (MTR) for 30 min (I) and the median fluorescence intensity (MFI) of MTR in GFP-positive cells was analyzed by flow cytometry (J). Based on the MTR MFI, the percentage of cells with decreased ΔΨm was calculated (K). Shown are the mean values ± SD of n = 2 biological replicates. Statistical differences were analyzed by one-way ANOVA and Tukey’s multiple comparison test. ** p ≤ 0.01. (L) Immunoblot analysis of Mfn2, Hsp60 (loading control) and GFP protein levels in GFP- or GFP-Lpg2444 expressing HEK293 FcγR cells treated with DMSO or 10 μM CCCP for 6h. Numbers indicated the fold change in Mfn2 levels normalized to the DMSO-treated GFP control. (M) AP-MS analysis workflow. HEK293T/17 cells were transfected with GFP or GFP-Lpg2444 in triplicate and lysed. GFP/GFP-Lpg2444 and their interaction partners (PPIs) were purified using GFP-Tap beads. Eluates were then subjected to LC-MS/MS to identify bound PPIs. (N) High-confidence PPI network of GFP-Lpg2444 (Bonferroni corrected false discovery rate < 5%). IMM proteins are highlighted in purple. (O) *Log_2_FC(WT/ΔdotA)* values of significantly regulated (adj.-p-value ≤ 0.05, |Log_2_FC| ≥ 1) ubiquitination (purple) and phosphorylation (grey) sites of the Lpg2444 PPIs SLC25A11, SLC25A5 and ATAD3A.

Strikingly, when compared to GFP-transfected cells, GFP-Lpg2444 expression induced the mitochondrial network to fuse into a single continuous structure (**Figures 6B** and **6G**, upper panel), suggesting that the effector might modulate mitochondrial dynamics. We therefore tested whether expression of the effector can rescue the disruption of the mitochondrial network upon exposure to the mitochondrial uncoupler carbonyl cyanide m-chlorophenyl hydrazone (CCCP). CCCP inhibits the oxidative phosphorylation machinery of mitochondria, thereby reducing the mitochondrial membrane potential (ΔΨm), which leads to mitochondrial fragmentation and mitophagy (Ishihara et al., 2003). As expected, CCCP led to the fragmentation of mitochondria in about 90% of GFP-expressing control cells (**Figures 6F, 6H** and **S6B**). In contrast, GFP-Lpg2444 expressing cells were refractory to mitochondrial fragmentation upon CCCP treatment (**Figures 6G-H** and **S6B**), indicating that the effector exerts functions that maintain mitochondrial network integrity. To test if GFP-Lpg2444 protects the cells from CCCP-induced ΔΨm dissipation, we treated GFP- or GFP-Lpg2444 expressing cells with DMSO or CCCP and measured the uptake of MitoTracker Red FM (MTR), a carbocyanide-based dye that accumulates in active mitochondria dependent on the mitochondrial membrane potential (**Figure 6I**), by flow cytometry. While CCCP led to a robust ΔΨm dissipation in GFP-transfected control cells, GFP-Lpg2444 transfected cells experienced a significantly lower decrease in ΔΨm upon CCCP treatment (**Figures 6J-K**), further supporting our hypothesis that Lpg2444 protects mitochondrial integrity. It has been proposed that CCCP-induced mitochondrial fragmentation and subsequent degradation via mitophagy requires the disruption of mitochondria-ER tethering (McLelland et al., 2018). This is mediated by the mitochondrial kinase PINK1 and the cytosolic E3 ubiquitin ligase Parkin which induce the phosphoubiquitination of mitofusin-2 (Mfn2), an OMM-located large GTPase that is required for mitochondrial fusion and ER-mitochondria tethering. This modification results in the proteasomal degradation of Mfn2 and release of mitochondria from the ER (McLelland et al., 2018; Tanaka et al., 2010). We therefore analyzed the levels of Mfn2 in GFP- or GFP-Lpg2444 expressing cells treated with DMSO or CCCP. Remarkably, while CCCP treatment led to a 50% reduction in Mfn2 protein levels GFP-expressing cells, Mfn2 was completely stabilized in GFP-Lpg2444 expressing cells (**Figure 6L**), indicating that Lpg2444 prevents mitochondrial damage that would lead to PINK1/Parkin activation. This further suggests that Lpg2444 has a mitoprotective role during stress conditions.

When we compared the amino acid sequences of Lpg2444 in the *L.p. Philadelphia* strain (this study) and its homologs in four other *L.p.* strains *(Lens, Corby, Alcoy, and Paris),* we found a high degree of conservation (**Figure S6C**). However, Lpg2444 (*L.p. Philadelphia* strain) had a unique N-terminal stretch of 21 amino acids that was entirely missing in the other strains. It is therefore possible that the Lpg2444 homologs might not be targeted to host cell mitochondria. If this additional sequence is required for mitochondrial targeting of the effector remains to be established. The amino acid sequence of Lpg2444 does not share similarity with any known prokaryotic or eukaryotic proteins or protein domains. Protein structure prediction with RaptorX (Källberg et al., 2012) suggested that Lpg2444 is a helical repeat protein (60% alpha-helices, 39% loops, p-value 2.37e-03) (**Figure S6D**). Prediction of transmembrane domains with TMPred (Hofmann and Stoffel, 1993) revealed a putative C-terminal membrane anchor (**Figure S6E**). This prediction might hint towards a protein scaffolding function of Lpg2444. We therefore set out to identify potential protein-protein interactions (PPIs) of Lpg2444 in host cells. To this end, we expressed the GFP-tagged Lpg2444 effector, GFP or the empty vector alone in HEK293T cells, enriched the baits from cell lysates with a GFP-Trap affinity resin and subjected them to affinity purification-mass spectrometry (AP-MS) (**Figure 6M**). A small aliquot was loaded on a polyacrylamide-gel and proteins were visualized with silver stain (**Figure S6F**). After background subtraction, we identified 15 Lpg2444 high confidence interactions (Bonferroni corrected false discovery rate < 5%) and analyzed these interaction partners via STRING (Jensen et al., 2009) (**Figure 6N** and **Table S5**). Strikingly, the PPIs for Lpg2444 included five IMM proteins, supporting our subcellular localization data and further suggesting that Lpg2444 localizes to or has contact with the IMM. Interestingly, three of these IMM interaction partners were also identified in our global proteomics analysis and showed increased phosphorylation (SLC25A11, ATAD3A) or de-ubiquitination (SLC25A5) upon infection with the *L.p.* WT strain at both 1 h and 8 h p.i., highlighting a potential role of these proteins during infection (**Figure 6O**). SLC25A5/ANT2 is of special interest as it has been shown to mediate ADP/ATP transport across the IMM and to be a component of the mitochondrial permeability transition pore (Halestrap and Davidson, 1990; Karch et al., 2019). Furthermore, SLC25A5/ANT2 has recently been implicated in the regulation of mitophagy (Hoshino et al., 2019). It will be highly interesting to functionally characterize the interaction between Lpg2444 and SLC25A5/ANT2 and to establish its role during *L.p.* infection.

## DISCUSSION

### Dynamic proteomics approach to identify novel host cell targets of *L.p*

In this study, we used a global proteomics approach to identify novel host cell signaling pathways, organelles and proteins that are regulated during *L.p.* infection through the secretion of bacterial effectors. To our knowledge, this is the most comprehensive resource of the regulation and/or modification of host cell proteins upon *L.p.* infection to date. Our analysis does not only include the quantification of changes in the host cell proteome, but also provides a spatiotemporal analysis of these changes to highlight the dynamic regulation of different predicted host cell organelles and signaling pathways in response to infection. This analysis showed expected hits, but also revealed novel exciting leads. For example, we found that peroxisomal proteins are regulated during *L.p.* infection, suggesting a previously unexplored role of peroxisomes during this process. Furthermore, we show for the first time that the mitochondrial proteome is reshaped during *L.p.* infection in a T4SS-dependent manner. The involvement of bacterial effectors could be either direct, and therefore lead to the identification of novel functions and targets of *L.p.* effectors, or indirect, giving insights on how the cell responds to the perturbation of homeostatic pathways mediated by the combined actions of the effectors. Importantly, the regulation of protein ubiquitination and phosphorylation reported in this study might include modifications mediated by *L.p.* kinases/phosphatases or ubiquitin ligases/deubiquitinating enzymes and can thereby lead to the identification of novel targets of these effectors.

### *L.p.* induces a mitochondrial stress response

Mitochondria are vital organelles with diverse functions, including energy generation and metabolism, calcium signaling, lipid and amino acid metabolism, and apoptotic signaling (Lebeau et al., 2018). It is therefore not surprising that many intracellular pathogens target mitochondria to benefit or protect themselves from these functions (Tiku et al., 2020). Although the recruitment of mitochondria to the LCV has been described almost four decades ago (Horwitz, 1983), the functional consequences of this interaction and the role of mitochondria during *L.p.* infection remain largely elusive. A recent report suggests that the *L.p.* effector MitF modulates mitochondrial dynamics to shift mitochondrial metabolism towards a Warburg-like metabolism in macrophages, thereby promoting bacterial replication (Escoll et al., 2017). Other groups have reported that certain *L.p.* effectors can translocate into host cell mitochondria to modulate lipid metabolism and metabolite transport (Degtyar et al., 2009; Dolezal et al., 2012). However, further research remains to be done to understand the manipulation of host cell mitochondria by *L.p.* in more detail. Our data reveal for the first time that the mitochondrial proteostasis network is perturbed and regulated during *L.p.* infection in a T4SS-dependent manner. This response shares similarities with the UPR^mt^, a homeostatic stress signaling response induced upon accumulation of misfolded proteins in the mitochondrial matrix (Shpilka and Haynes, 2017). Interestingly, is has been shown that the UPR^mt^ is induced as an adaptive response in cells experiencing the Warburg effect (Kenny et al., 2019), suggesting that the previously reported effector-induced shift in mitochondrial metabolism (Escoll et al., 2017) might cause the induction of a UPR^mt^-like stress response in infected cells. However, it is also possible that the translocation of bacterial effectors into host cell mitochondria (Degtyar et al., 2009; Dolezal et al., 2012) might activate this stress response. Furthermore, the effector-mediated inhibition of host cell translation during *L.p.* infection might contribute to the induction of a mitochondrial stress response (Andréasson et al., 2019). Much to our surprise, we found that the induction of classic ISR-activated target genes, such as *CHOP, GADD34* and *ATF3,* upon *L.p.* infection does not require signaling through the ISR, suggesting an alternative signaling pathway. This finding was interesting as it somewhat reflects the current conundrum in the field of mitochondrial stress research. While the ISR is clearly required for stress signaling during certain types of mitochondrial stress induced by small molecules such as CCCP or oligomycin (Guo et al., 2020; Quirós et al., 2017), it’s contribution during other forms of mitochondrial stress (e.g. upon treatment with CDDO or GTPP) has been unclear (Fessler et al., 2020; Münch and Harper, 2016). For instance, although tunicamycin, a potent inducer of ER stress, the mitochondrial uncoupler CCCP and the UPR^mt^ inducer GTPP all activate CHOP, a central component of the ISR, only GTPP treatment leads to the upregulation of mitochondrial chaperonins, and the up-regulation of CHOP upon GTPP treatment does not seem to require any of the ISR kinases, further supporting the existence of stressor-specific ISR signatures and signaling routes (Münch and Harper, 2016). As all these studies rely on the use of small molecule inducers of mitochondrial stress and might induce unspecific effects in the cell, the use of *L.p.* may offer a novel, alternative tool to study mitochondrial stress responses in the context of bacterial infection. Our study revealed the selective and strong induction of the protein ATF3 upon *L.p.*-induced stress, while ATF4 and CHOP translation was suppressed. This might highlight an underappreciated role of ATF3 in mitochondrial stress signaling, which has also been proposed independently of our study (Münch and Harper, 2016), but has remained unexplored so far. We are currently addressing this question in our laboratory.

### Lpg2444 is a novel mitochondrially targeted *L.p.* effector with mitoprotective functions

The notion that *L.p.* induces mitochondrial stress in an T4SS-dependent manner led us to search for novel *L.p.* effectors targeting host cell mitochondria. We show that Lpg2444, a previously uncharacterized effector with predicted alpha-helical structure, localizes to host cell mitochondria upon ectopic expression in cells. Strikingly, the effector protected mitochondria from CCCP-induced ΔΨm dissipation and mitochondrial fragmentation and interacted with SLC25A5/ANT2, an IMM protein involved in mitophagy (Hoshino et al., 2019). These findings suggest a mitoprotective function of Lpg2444, especially during conditions of mitochondrial stress. During the course of infection, Lpg2444 might therefore counteract insults to mitochondria that are mediated by other *L.p.* effectors, such as MitF (Escoll et al., 2017), to prevent premature mitochondrial apoptosis of the host cell. In line with this hypothesis, it has been reported that the effectors SdhA and SidF prevent cell death by inhibiting mitochondrial fission and cytochrome c release and by binding to and inhibiting the pro-apoptotic factor Bcl2-rambo, respectively (Banga et al., 2007; Laguna et al., 2006). Our lab is currently defining the mechanism by which Lpg2444 exerts its mitoprotective functions and the role that the effector plays in the context of *L.p.* infection.

## ACKNOWLEDGEM ENTS

We thank Dr. Philipp Schlaermann for preparation of cell pellets for the proteomics experiment and Dr. Lan Wang for help with the structure prediction of Lpg2444. We thank Dr. Peter Walter for scientific advice. We thank Dr. Advait Subramanian for critically reading the manuscript. S.M. is supported by the National Institutes of Health RO1 grant AI118974 and an award from the Pew Charitable Trust (A129837).

## AUTHOR CONTRIBUTIONS

Conceptualization, S.M., J.N., D.J-M., D.L.S., and N.J.K.; Methodology, S.M., J.N., E.S., G.J., D.J-M., D.L.S., and N.J.K.; Data Curation, D.J-M., D.L.S., and J.N.; Validation, J.N.; Formal Analysis, D.J-M. and J.N.; Investigation, J.N., T.M.; Writing – Original Draft, J.N. and S.M.; Writing – Review & Editing, J.N., S.M., D.L.S., and D.J-M.; Visualization, J.N, D.J.-M..; Funding Acquisition, S.M., D.L.S., and N.J.K.; Resources, S.M., D.L.S., and N.J.K.; Supervision, S.M., D.L.S., and N.J.K..

## METHODS

### LEAD CONTACT AND MATERIALS AVAILABILITY

Further information and requests for resources and reagents should be directed to and will be fulfilled by the Lead Contact, Shaeri Mukherjee. This study did not generate new unique reagents.

### EXPERIMENTAL MODEL AND SUBJECT DETAILS

#### Cell lines

HEK293T cells (female), HEK293 cells (female) stably expressing the Fcγ receptor III (HEK293 FcγR cells) and HeLa cells (female) stably expressing the Fcγ receptor III (HeLa FcγR cells) (Gifts from the lab of Dr. Craig Roy at Yale University) were cultured in Dulbecco’s Modified Eagle’s Medium (DMEM, GIBCO) containing 10% fetal bovine serum (FBS, VWR) at 37°C and 5% CO_2_.

#### Bacterial Strains

The *L.p.* strains LP01 WT and LP01 *ΔdotA* (Gifts from the lab of Dr. Craig Roy at Yale University) were cultivated on Charcoal Yeast Extract (CYE) agar plates or ACES Yeast Extract (AYE) medium.

### METHOD DETAILS

#### Infection of cultured mammalian cells with *L.p*

Infections with *L.p.* were performed as previously described (Treacy-Abarca and Mukherjee, 2015). *L.p.* heavy patches grown for 48 h on CYE plates were used for overnight liquid cultures in AYE medium supplemented with 0.33 mM Fe(NO_3_)_2_ and 3.3 mM L-cysteine until reaching an OD600 of ~3. *L.p.* from the overnight culture was enumerated and the appropriate amount was opsonized with *L.p*.-specific antibodies at a dilution of 1:2000 in cell growth medium for 20 min. HEK293 FcγR were grown on poly-lysine coated cell culture plates to a confluency of 80% and infected with the *L.p.* WT strain or the isogenic *ΔdotA* mutant strain at an MOI of 1-100 as indicated. The infection was synchronized by centrifugation of the plates at 500xg for 5 min. Cells were washed three times with warm PBS after 1 h of infection and fresh growth medium was added. Cells were collected for down-stream processing at the indicated time-points.

#### Sample preparation for proteomics analysis

Uninfected HEK293 FcγR cells or HEK293 FcγR infected for 1 h or 8 h with the *L.p.* WT strain or the isogenic *ΔdotA* mutant were infected at an MOI of 100. Cells were washed with ice-cold PBS, collected and the pellet was frozen at −80°C. Cell pellets were lysed by probe sonication in three pulses of 20% amplitude for 15 s in a lysis buffer consisting of: 8 M urea, 150 mM NaCl, 100 mM ammonium bicarbonate, pH 8; added per 10 ml of buffer: 1 tablet of Roche mini-complete protease inhibitor EDTA free and 1 tablet of Roche PhosSTOP. In order to remove insoluble precipitate, lysates were centrifuged at 16,100 g at 4°C for 30 min. A Bradford Assay (Thermo) was performed to measure protein concentration in cell lysate supernatants. 6 mg of each clarified lysate was reduced with 4 mM tris(2-carboxyethyl)phosphine for 30 min at room temperature and alkylated with 10 mM iodoacetamide for 30 min at room temperature in the dark. Remaining alkylated agent was quenched with 10 mM 1,4-dithiothreitol for 30 min at room temperature in the dark. The samples were diluted with three starting volumes of 100 mM ammonium bicarbonate, pH 8.0, to reduce the urea concentration to 2 M. Samples were incubated with 50 μg of sequencing grade modified trypsin (Promega) and incubated at room temperature with rotation for 18 hr. The sample pH was reduced to approximately 2.0 by the addition of 10% trifluoroacetic acid (TFA) to a final concentration of 0.3% trifluoroacetic acid. Insoluble material was removed by centrifugation at 16,000 g for 10 min. Peptides were desalted using SepPak C18 solid-phase extraction cartridges (Waters). The columns were activated with 1 ml of 80% acetonitrile (ACN), 0.1% TFA, and equilibrated 3 times with 1 ml of 0.1% TFA. Peptide samples were applied to the columns, and the columns were washed 3 times with 1 ml of 0.1% TFA. Peptides were eluted with 1.2 ml of 50% ACN, 0.25% formic acid. Peptides were divided for global protein analysis (10 μg), phosphopeptide enrichment (1 mg), or diGly-enrichment (remaining sample), and lyophilized.

#### Phosphopeptide enrichment by immobilized metal affinity chromatography

Iron nitriloacetic acid (NTA) resin were prepared in-house by stripping metal ions from nickel nitroloacetic acid agarose resin with 100 mM ethylenediaminetetraacetic acid, pH 8.0 four times. Resin was washed twice with water and 100 mM iron(III) chloride was applied four times. The iron-NTA resin was washed twice with water and once with 0.5% formic acid. Iron-NTA beads were resuspended in water to create a 25% resin slurry. 60 μl of Fe-NTA resin slurry was transferred to individual Silica C18 MicroSpin columns (The Nest Group) pre-equilibrated with 100 μl of 80% CAN, 0.1% TFA on a vacuum manifold. Subsequent steps were performed with the Fe-NTA resin loaded above the Silica C18 columns. Dry peptide samples were resuspended in a solution of 200 μl 75% ACN 0.15% TFA. Peptide samples were mixed twice with the Fe-NTA resin, allowing the peptides to incubate for 2 minutes between each mixing step. The resin was rinsed four times with 200 μl of 80% ACN, 0.1% TFA. In order to equilibrate the columns, 200 μl of 0.5% formic acid was applied twice to the resin and columns. Peptides were eluted from the resin onto the C18 column by mixing and incubating the Fe-NTA resin with 200 μl of 500 mM potassium phosphate, pH 7.0 for 2 minutes. The elution step was repeated once. Peptides bound to the C18 column were washed three times with 200 μl of 0.5% formic acid. The C18 columns were removed from the vacuum manifold and eluted twice by centrifugation at 1000g with 75 μl of 50% ACN, 0.25% formic acid. Peptides were dried with a centrifugal adaptor and stored at −20°C until analysis by liquid chromatograph and mass spectrometry.

#### Di-glycine peptide enrichment by immunoprecipitation

Peptide samples were subjected to ubiquitin remnant immunoaffinity. 10 uL of PTMScan^®^ Ubiquitin Remnant Motif (K-ε-GG) Antibody Bead Conjugate purification (Cell Signaling) slurry was used per 1 mg peptide sample. Ubiquitin remnant beads were washed twice with IAP buffer, then split into individual 1.7 mL low bind tubes (Eppendorf) for binding with peptides. Peptides were dried with a centrifugal evaporator for 12 hours to remove TFA in the elution. The lyophilized peptides were resuspended in 1 ml of IAP buffer (50 mM 4-morpholinepropnesulfonic acid, 10 mM disodium hydrogen phosphate, 50 mM sodium chloride, pH 7.5). Peptides were sonicated and centrifuged for 5 minutes at 16,100g. The soluble peptide supernatant was incubated with the beads at 4°C for 90 minutes with rotation. Unbound peptides were separated from the beads after centrifugation at 700g for 60 seconds. Beads containing peptides with di-glycine remnants were washed twice with 500 μL of IAP buffer, then washed twice with 500 μL of water, with a 700g 60s centrifugation to allow the collection of each wash step. Peptides were eluted twice with 60 μL of 0.15% TFA. Di-glycine remnant peptides were desalted with UltraMicroSpin C18 column (The Nest Group). Desalted peptides were dried with a centrifugal adaptor and stored at −20°C until analysis by liquid chromatograph and mass spectrometry.

#### Mass spectrometry data acquisition and analysis

Samples were resuspended in 4% formic acid, 4% acetonitrile solution, separated by a reversed-phase gradient over a nanoflow column (360 μm O.D. x 75 μm I.D.) packed with 25 cm of 1.8 μm Reprosil C18 particles with (Dr. Maisch), and directly injected into an Orbitrap Fusion Lumos Tribrid Mass Spectrometer (Thermo). Total acquisition times were 120 min for protein abundance, 100 min for phosphorylation, and 70 min for ubiquitylation analyses. Specific data acquisition settings are detailed in **Table S6**. Raw MS data were searched with MaxQuant against both the human proteome (UniProt canonical protein sequences downloaded January 11, 2016) and the *Legionella Pneumophila Philadelphia* proteome (downloaded July 17, 2017). Peptides, proteins, and PTMs were filtered to 1% false discovery rate in MaxQuant (Cox and Mann, 2008).

#### Functional enrichment and network analysis

For **Figure 1K**, a list of all significantly regulated proteins (adj.-p-value ≤ 0.05, |Log_2_FC| ≥ 1) in WT *vs. ΔdotA*-infected cells in the AB dataset or in the combined UB/PH datasets was analyzed for significantly overrepresented gene ontology (GO) terms (Biological Processes) and biological pathways (KEGG, Reactome) with the g:Profiler g:GO St tool (Raudvere et al., 2019). The g:SCS algorithm was selected for multiple testing correction. The *Homo sapiens* genome (only annotated genes) was selected as the background gene list. For **Figure 2D**, a list of all kinases with predicted up-regulated activity in WT *vs. ΔdotA*-infected cells was analyzed for significantly overrepresented biological pathways (KEGG) and gene ontology (GO) terms (Biological Processes) using the stringApp in Cytoscape (Doncheva et al., 2019; Shannon et al., 2003). The false-discovery rate (FDR) was used for multiple testing correction. The *Homo sapiens* genome (only annotated genes) was used as the background gene list. For **Figures 4B** and **S4A**, a list of all significantly regulated proteins/proteoforms (adj.-p-value ≤ 0.05, ILog_2_FC| ≥ 1) in WT *vs. ΔdotA*-infected cells (AB, UB and PH datasets combined) was analyzed for significantly overrepresented gene ontology (GO) terms, biological pathways, INTERPRO Protein Domains and Features and UniProt Keywords using the stringApp in Cytoscape (Doncheva et al., 2019; Shannon et al., 2003). The FDR was used for multiple testing correction. The *Homo sapiens* genome (only annotated genes) was used as the background gene list.

#### Prediction of kinase activity and complex regulation

Kinase activities and complex regulation were inferred from all significantly regulated phosphosites (adj.-p-value ≤ 0.05, |Log_2_FC| ≥ 1) in WT *vs. ΔdotA*-infected using PhosFate Profiler (Ochoa et al., 2016). The inferred kinase activities (branch color), p-values (node color) and substrate numbers (node size) were mapped on the human kinome tree with CORAL (Metz et al., 2018) for data visualization.

#### Subcellular mapping of proteomics data

For **Figures 3A-B**, the number of significantly regulated proteins (adj.-p-value ≤ 0.05, |Log_2_FC| ≥ 1) in WT *vs. ΔdotA*-infected cells (AB, UB and PH) in each subcellular compartment was quantified based on their primary ECO or, if not available, on their documented subcellular localization in The Human Protein Atlas database (http://www.proteinatlas.org) (Thul et al., 2017).. The range of regulated proteins was assigned to a color range and mapped on the host cell using a custom-designed template in Adobe Illustrator. For **Figures 3C** and **S3**, a list of significantly regulated proteins (adj.-p-value ≤ 0.01, |Log_2_FC| ≥ 1 (**S3**) or 2 (**3C**)) in WT *vs. ΔdotA*-infected cells (AB, UB and PH) with their primary subcellular annotation was imported into Cytoscape (Doncheva et al., 2019; Shannon et al., 2003) and proteins were colored according to their Log_2_FC value. The outline of each represented protein reflects the regulated dataset (circle: AB, octagon: UB, rounded square: PH). The resulting graphics were then imported into custom-designed templates in Adobe Illustrator.

#### Cell lysis and immunoblot analysis

HEK293 FcγR cells grown on poly-lysine coated plates were treated as indicated, washed three times with ice-cold PBS and harvested with a cell scraper. The cell pellets were frozen at −80°C. Cell pellets were resuspended in RIPA buffer supplemented with cOmplete Protease Inhibitor Cocktail (Roche) and PhosSTOP (Roche) and lysed under constant agitation for 30 min at 4°C. Cell debris was removed by centrifugation at 12,000xg for 20 min at 4°C. Protein concentration was measured using the DC Protein Assay (Bio-Rad) or the Pierce™ BCA Protein Assay Kit (Thermo Fisher Scientific). For each sample, 20-30 μg of proteins were denatured in SDS sample buffer/5% β-mercaptoethanol at 95°C for 5 min, loaded on 8-12% SDS-polyacrylamide gels and separated by SDS-PAGE. Proteins were transferred to PVDF membranes (0.45 μm, Millipore) at 30 V, 4°C for 16 h. Membranes were washed with PBS-T (PBS/ 0.1% Tween-20 (Thermo Fisher Scientific)), blocked with 5% Blotting Grade Blocker Non Fat Dry Milk (Bio-Rad) for 1 h at room temperature and incubated with the primary antibodies diluted in blocking buffer/0.02% (w/v) sodium azide overnight at 4°C. The primary antibodies were diluted as follows: ATF3 1:1000 (Cell Signaling Technology), TIMM17A 1:1000 (Proteintech), OPA-1 1:1000 (Cell Signaling Technology), MRPP3 1:1000 (Proteintech), TIMM23 1:1000 (Proteintech), CHOP 1:500 (Cell Signaling Technology), ATF4 1:1000 (Cell Signaling Technology), HSP60 1:20000 (Proteintech), GFP 1:2000 (Proteintech), alpha-Tubulin 1:5000 (Proteintech), Mfn2 1:1000 (Proteintech), Phospho-c-Jun (Ser73) 1:1000 (Cell Signaling Technology). Membranes were washed three times with PBS-T and incubated with Goat Anti-Mouse IgG (H+L) HRP Conjugate (Thermo Fisher Scientific), Goat Anti-Rabbit IgG (H+L) (Thermo Fisher Scientific), HRP Conjugate, or Protein A HRP conjugate (Cell Signaling Technology) diluted at 1:5000 in blocking buffer for 45 min at room temperature. After three washes with PBS-T, membranes were incubated with Amersham ECL Western Blotting Detection Reagent (Global Life Science Solutions) for 1 min and imaged on a ChemiDoc Imaging System (BioRad).

#### RT-qPCR

HEK293 FcγR cells were grown on poly-lysine coated 6well plates or 350 mm dishes and treated as indicated. After treatment, the medium was removed, and cells were lysed in TRIzol reagent for 5 min at room temperature. RNA was extracted using the Direct-zol RNA Miniprep Plus kit (Zymo Research) according to the manufacturer’s instructions. For each sample, 500 ng RNA were used for cDNA synthesis with the SuperScript III kit (Invitrogen) using oligo(dT)20 primers. Transcript levels were analyzed by qPCR using the BioRad iTaq SYBR Green kit. For each reaction, 5 ng of cDNA was used. Three technical replicates for each sample were analyzed on the BioRad CFX384 Touch™ Real-Time PCR Detection System. Data were processed using the ΔΔCt method.

#### Immunofluorescence

HEK293 FcγR or HeLa FcγR cells were grown on poly-lysine coated coverslips in 24well cell culture plates. Cells were treated as indicated, washed three times with PBS and fixed in 4% paraformaldehyde/PBS for 15 min at room temperature. All further steps were done at room temperature as well. For **Figures S6A**, cells were permeabilized with 0.1% Triton X-100/PBS for 20 min followed by blocking with 1% BSA/PBS for 1h. Cells were stained with primary antibodies diluted in 1% BSA/PBS for 1h, washed three times with PBS and stained with secondary antibodies diluted in 0.2% BSA/PBS for 45 min. For **Figures 6B, 6F-G** and **S6B**, cells were permeabilized and blocked with 0.5% saponin/2% BSA/PBS for 1h, stained with primary antibodies diluted in 0.5% saponin/2% BSA/PBS for 1h, washed three times with PBS and stained with secondary antibodies diluted in 0.5% saponin/2% BSA/PBS for 45 min. Cells were then stained with Hoechst33342 at 1:2000 in PBS for 10 min and washed three times with PBS. Coverslips were dipped three times into purified ddH_2_O to remove salts, dried and mounted on microscopy glass slides with Prolong Diamond antifade 5 (Thermo Fisher Scientific). Slides were cured overnight at room temperature and imaged the next day on a spinning disk Eclipse Ti2-E inverted microscope (Nikon). The following antibody dilutions were used: primary antibodies: ATF3 1:100 (Cell Signaling Technology), *L.p.* 1:2000 (Thermo Fisher Scientific), TOM20 1:200 (BD Biosciences); secondary antibodies: Goat anti-Rabbit IgG (H+L) Highly Cross-Adsorbed Secondary Antibody, Goat anti-Mouse IgG (H+L) Highly Cross-Adsorbed Secondary Antibody, Alexa Fluor 633 1:500 (Thermo Fisher Scientific) or Alexa Fluor 488 1:500 (Thermo Fisher Scientific).

#### Generation of GFP-tagged *L.p.* effectors

*L.p.* effectors cloned into the pMSCV-6X Myc backbone (gift from the laboratory of Dr. Russel Vance at University of California, Berkeley) were PCR-amplified using the primers pEGFP_attB1_F: 5’-CTGTACAAGTCCGGCCGGACTCAAAGTTTGTACAAAAAAGCAGGCTTC-3’ and pEGFP_attB2_R: 5’-AGTTATCTAGATCCGGTGGATCGACCACTTTGTACAAGAAAGCTGG-3’. The pEGFP-C2 mammalian expression vector was cut with the restriction enzymes BglII and BamH1. The amplified *L.p.* effectors were then inserted into the vector using the Gibson Assembly Master Mix (NEB).

#### Cell transfections

All transfections were performed with jetPRIME (Polyplus). HEK293 FcyR or HeLa FcyR cells were grown to 60% confluency and transfected according to the manufacturer’s recommendations. For transfection of plasmid DNA, 0.25 μg DNA was used for 24well plates, 1 μg DNA for 60 mm plates and 10 μg DNA for 150 mm plates. 12h after transfection, cells were treated as indicated and analyzed or harvested. For siRNA transfections, 42 pmoles of siRNA (MISSION^®^ esiRNA targeting human YME1L1, #EHU115921, or MISSION^®^ siRNA Universal Negative Control #1, #SIC001, MilliporeSigma) were used for 60 mm plates. 24h after transfection, cells were treated as indicated and harvested.

#### Subcellular fractionation

For analysis of cytosolic and mitochondrial fractions, HEK293 FcγR cells were grown on 150 mm cell culture plates (two plates for each condition) to 60% confluency and transfected with GFP or GFP-Lpg2444 plasmids. 18h after transfection, cells from two plates were pooled in 10 ml icecold PBS and collected by centrifugation at 850xg for 5 min at 4°C. Cells were resuspended in 1.5 ml ice-cold PBS, transferred in a 2 ml microcentrifuge tubes and collected by centrifugation. From this cell pellet, the isolation of mitochondria and cytosol was performed with the Mitochondria Isolation Kit for Cultured Cells (Thermo Scientific) according to the manufacturer’s instructions. Mitochondrial pellets were resuspended in 100 μl RIPA buffer supplemented with EDTA-free protease inhibitor cocktail (EMD Millipore) under constant agitation for 30 min at 4°C. Mitochondrial and cytosolic fractions were analyzed by immunoblot as described above.

#### Flow cytometry-based ΔΨ_m_ assay

HEK293 FcyR cells were grown on poly-lysine coated 60 mm cell culture dishes and transfected with GFP- or GFP-Lpg2444. 12h after transfection, cells were treated with 200 nM MitoTracker Red FM (Thermo Fisher Scientific) and DMSO or 20 μM CCCP for 30 min. Cells were trypsinized, collected with a cell scraper and washed once with PBS. Cell pellets were carefully resuspended in phenol-red free cell culture medium (GIBCO)/10% FBS (VWR) at 10^6^ cell/ml and analyzed on a BD FACSCalibur analyzer (BD Biosciences). The FSC/SSC channels were used to exclude cell debris from the analysis. The FL2 channel was used to gate for the GFP-positive cells. The FL4 channel was used to analyze the MitoTracker Red FM signal in GFP-positive cells. Untransfected cells stained with MitoTracker Red FM only and unstained GFP-transfected cells were used as compensation controls.

#### Affinity purification-mass spectrometry (AP-MS)

HEK293T cells were cultured in Dulbecco’s Modified Eagle’s Medium (Corning) supplemented with 10% Fetal Bovine Serum (Gibco, Life Technologies) and 1% Penicillin-Streptomycin (Corning) and maintained at 37°C in a humidified atmosphere of 5% CO_2_. For each immunoprecipitation, ten million HEK293T cells were plated per 15-cm dish and transfected with up to 15 μg of individual GFP-tagged expression constructs after 20-24 hours. Total plasmid was normalized to 15 μg with vector DNA and complexed with PolyJet Transfection Reagent (SignaGen Laboratories) at a 1:3 μg: μl ratio of plasmid to transfection reagent based on manufacturer’s recommendations. After 40 hours, cells were dissociated at room temperature using 10 ml Dulbecco’s Phosphate Buffered Saline without calcium and magnesium (D-PBS) supplemented with 10 mM EDTA for at least 5 minutes and subsequently washed with 10 ml D-PBS. These steps were followed by centrifugation at 200 xg, 4°C for 5 minutes. Cell pellets were frozen on dry ice and stored at −80°C. Three biological replicates were independently prepared for affinity purification. Cell pellets were thawed on ice for 15-20 minutes, suspended in 600 μl Lysis Buffer [IP Buffer (50 mM Tris-HCl, pH 7.4 at 4°C, 150 mM NaCl, 1 mM EDTA) supplemented with 0.5% Nonidet P 40 Substitute (NP40; Fluka Analytical) and cOmplete mini EDTA-free protease and PhosSTOP phosphatase inhibitor cocktails (Roche)], 1 mM DTT, and 75 U benzonase, and immediately frozen on dry ice for 10 minutes. Samples were partially thawed in a 37°C water bath and further incubated on a tube rotator for 15 minutes before centrifugation at 4,300 xg, 4°C for 15 minutes to pellet debris. The lysate was then added to 20 μl GFP-Trap beads (Chromotek) which were equilibrated twice with 1 ml Wash Buffer (IP Buffer supplemented with 0.05% NP40). After binding on a tube rotator for 2 hours, beads were washed one time with 1 ml Wash Buffer, followed by 2 times with Wash Buffer that was not supplemented with NP40. Proteins were eluted twice, each time with 50 μl 0.1 M Glycine pH 2.5 by gently agitating beads on a vortex mixer at room temperature for 10 minutes. After removing from beads, eluates were neutralized with 1 M Tris-HCl, pH 9.0 to adjust the solution pH to 8.0. To prepare samples for LC-MS/MS analysis, eluates were reduced by the addition of 1 mM DTT at 60°C for 15 minutes, cooled to room temperature, alkylated by the addition of 3 mM iodoacetamide for 45 minutes in the dark. Alkylation was quenched by the addition of 3 mM DTT and proteins were digested overnight at 37°C with 1 μg trypsin (0.5 μg/μl; Promega). Following digestion, peptides were acidified with trifluoroacetic acid (0.5% final, pH < 2), desalted using UltraMicroSpin Columns (PROTO 300 C18 300Å; The NEST Group) according to manufacturer’s specifications, and dried under vacuum centrifugation (CentriVap Concentrator, Labconco). Samples were resuspended in 4% formic acid, 4% acetonitrile solution, and separated by a 70-minute reversed-phase gradient over a nanoflow column (360 μm O.D. x 75 μm I.D.) packed with 25 cm of 1.8 μm Reprosil C18 particles (Dr. Maisch). Peptides were directly injected into an Orbitrap Fusion Tribrid Mass Spectrometer (Thermo), with all MS1 spectra collected in the orbitrap, and MS2 spectra collected in the ion trap. Raw MS data were searched with MaxQuant (Cox and Mann, 2008) against both the human proteome (Uniprot canonical protein sequences downloaded March 21,2018) and the *Legionella Pneumophila Philadelphia* proteome (downloaded July 17, 2017). Peptides and proteins were filtered to 1% false discovery rate in MaxQuant, and identified proteins were then subjected to protein-protein interaction scoring using SAINTexpress (Teo et al., 2014).

### QUANTIFICATION AND STATISTICAL ANALYSIS

Statistical analysis of quantifications obtained from MaxQuant was performed with the artMS Bioconductor package (version 0.9) (Jimenez-Morales et al., 2019). Each dataset (proteome phosphoproteome, and ubiquitinome) was analyzed independently. Quality control plots were generated using the artMS quality control functions. The site-specific relative quantification of posttranslational modifications required a preliminary step consisting of providing the ptm-site/peptide-specific annotation (“artmsProtein2SiteConversion()” function). artMS performs the relative quantification using the MSstats Bioconductor package (version 3.14.1) (Choi et al., 2014). Contaminants and decoy hits were removed. Samples were normalized across fractions by median-centering the Log_2_-transformed MS1 intensity distributions (**Fig. S1B**). Log_2_FC for protein/sites with missing values in one condition but found in >2 biological replicates of the other condition of any given comparison were estimated by imputing intensity values from the lowest observed MS1-intensity across samples peptides (Webb-Robertson et al., 2015); p-values were randomly assigned between 0.05 and 0.01 for illustration purposes. Statistically significant changes were selected by applying a Log_2_FC (>1.0 or <-1.0) and adjusted p-value (<0.05).

Statistical analysis of imaging, qPCR and flow cytometry data was performed with GraphPad Prism 8. Comparisons of data were performed by one-way ANOVA and Tukey’s multiple comparison test or by unpaired, two-tailed t-tests as indicated. p-values: ns p > 0.05, * p ≤ 0.05, p ≤ 0.01 **, p ≤ 0.001 ***, p ≤ 0.0001 ****.

## DATA AND CODE AVAILABILITY

The mass spectrometry data files (raw and search results) have been deposited to the ProteomeXchange Consortium (http://proteomecentral.proteomexchange.org) via the PRIDE partner repository with the dataset identifier PXD019217 (Vizcaino et al., 2016).

---

**Figure S1.**
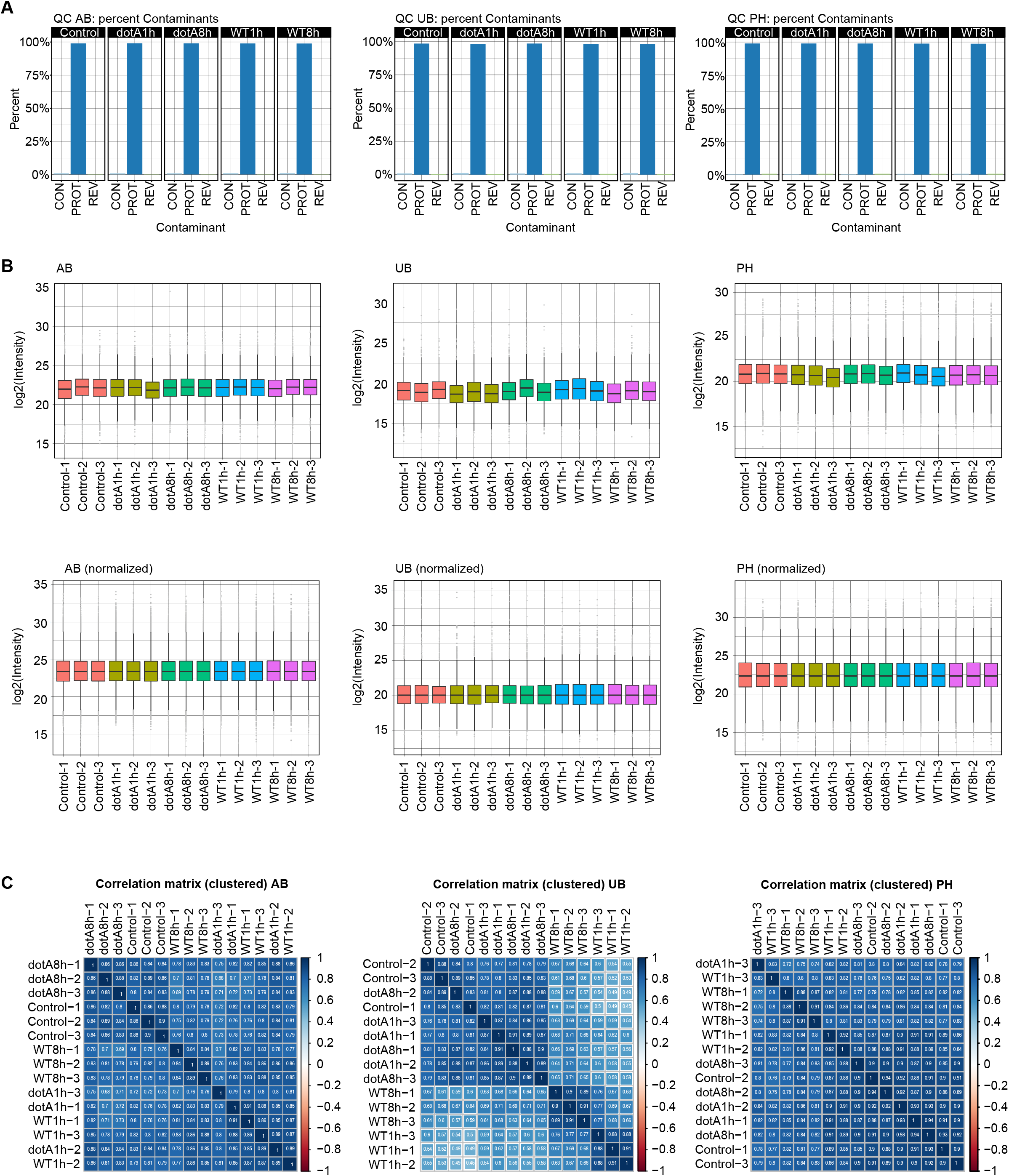
Quantification and quality control plots of proteomics data. Related to Figure 1. Quality control plots for each dataset (AB, UB, PH) were generated using the artMS Bioconductor package (version 0.9) (Jimenez-Morales et al., 2019). (A) Percent of contaminants (CON), proteins (PROT) and reversed sequences (REV) in each experimental condition (control, dotA-1h, dotA-8h, WT-1h, WT-8h) were quantified to adjust the false-discovery-rate (FDR). (B) Samples were normalized across fractions by median-centering the Log_2_-transformed MS1 intensity distributions. (C) Correlation matrices showing the clustering of the different experimental conditions.

**Figure S2.**
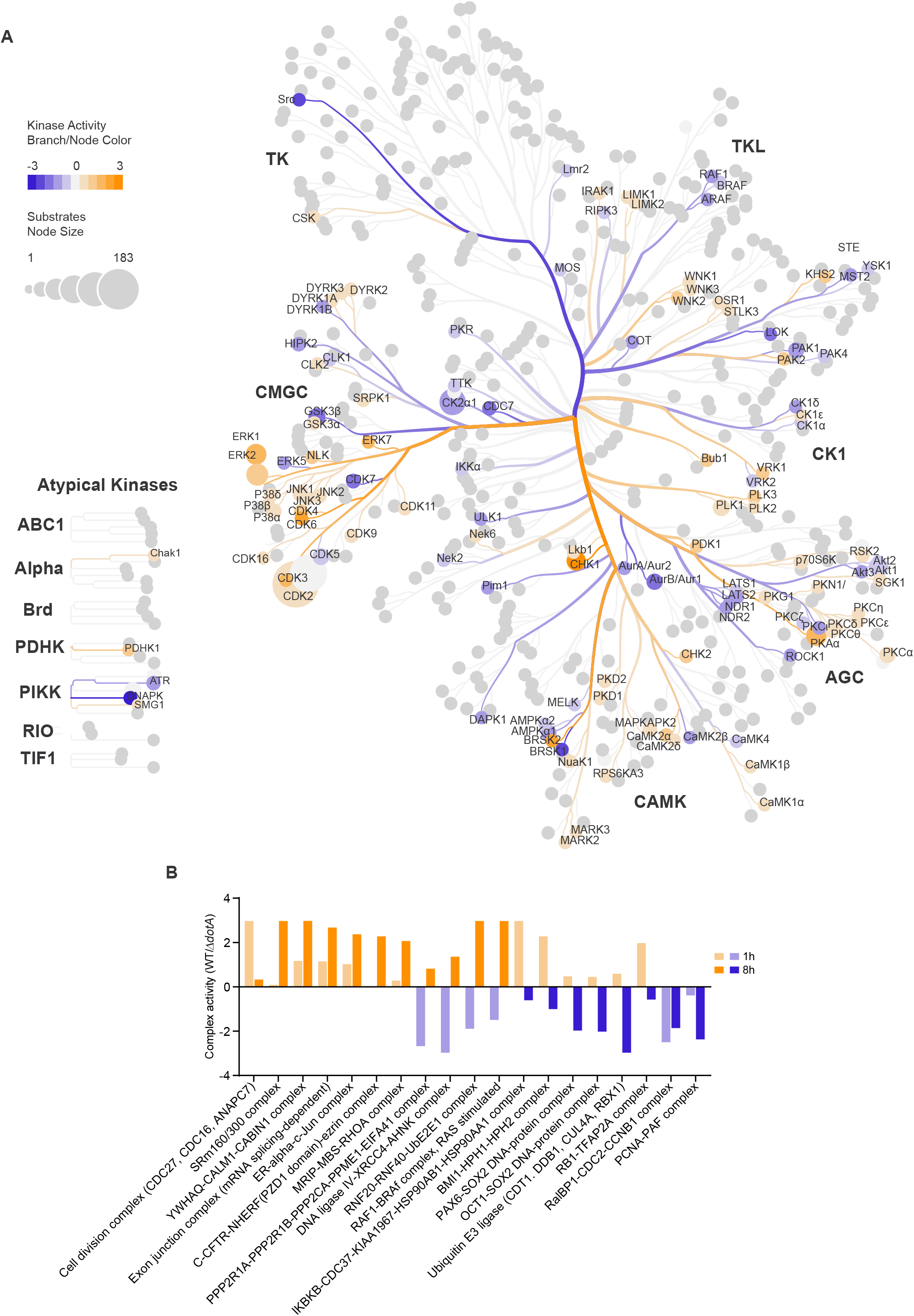
Prediction of complex regulation in *L.p.* infected cells. Related to Figure 2. (A) Host cell kinase activities in WT-*vs. ΔtdotA*-infected cells at 1h p.i. were inferred with PhosFate Profiler (Ochoa et al., 2016) based on regulated phosphorylated sites, and mapped on the kinase tree with CORAL (Metz et al., 2018). Kinase activity is indicated by the branch and node color, and the number of substrates by the node size. Names of kinase families and regulated kinases are highlighted. (B) Based on the prediction of kinase activities, complex activities were inferred with PhosFate Profiler (Ochoa et al., 2016). The bar graph shows the most significantly regulated complexes (p-value ≤ 0.01) in WT-*vs. ΔdotA*-infected cells.

**Figure S3.**
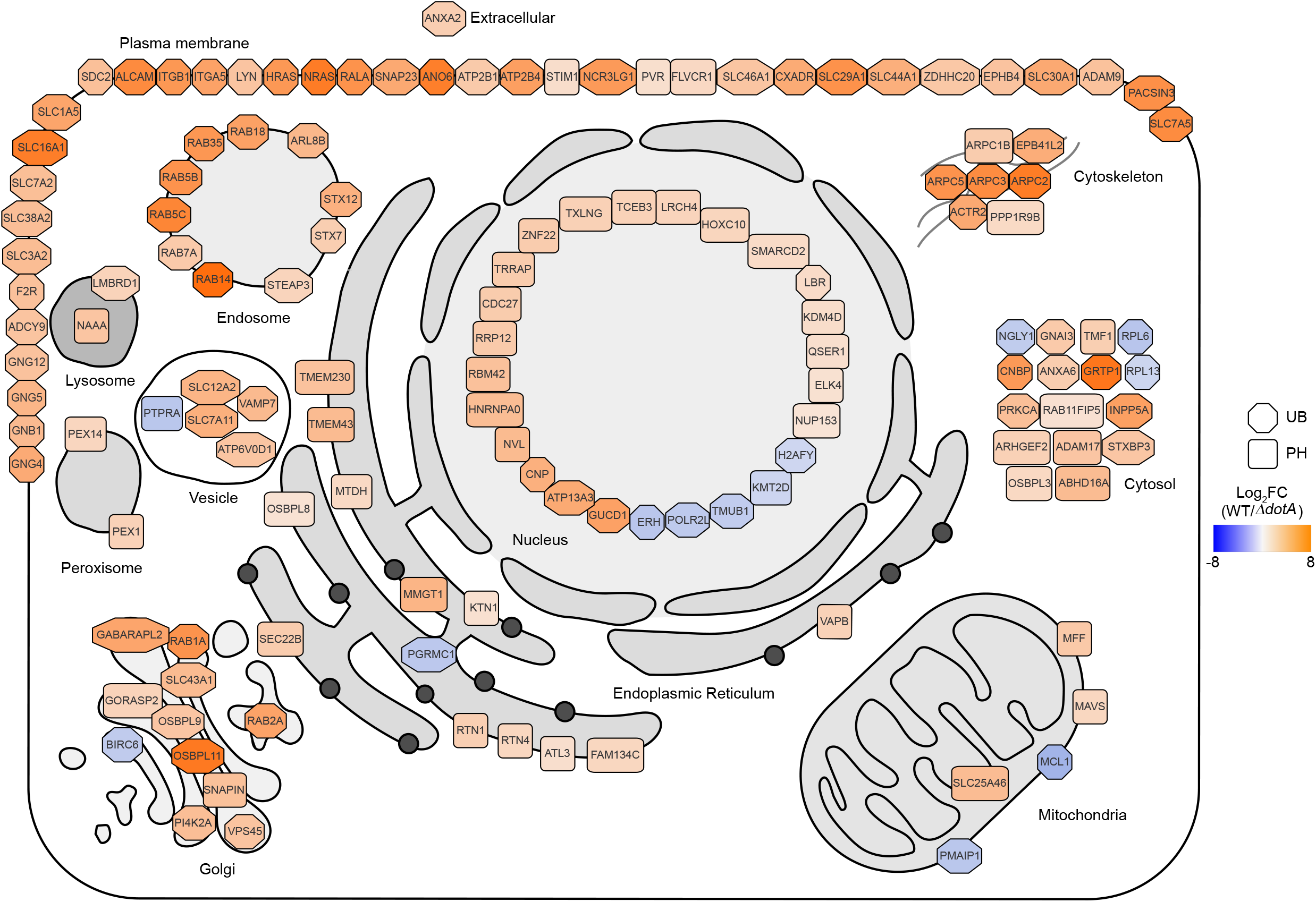
Effector-dependent spatiotemporal proteomic changes at 1h p.i.. Related to Figure 3. Highly significantly regulated proteins (adj. p-value ≤ 0.01, |Log_2_FC(WT/*ΔdotA*)| ≥ 1) were mapped on the host cell organelles according to their primary ECO. The Log_2_FC(WT/*ΔdotA*) values are indicated by a color scale (orange: up-regulated, blue: down-regulated), the dataset is indicated by the shape of the icon (octagon: UB, rounded square: PH). AB was not regulated at this significance cut-off.

**Figure S4.**
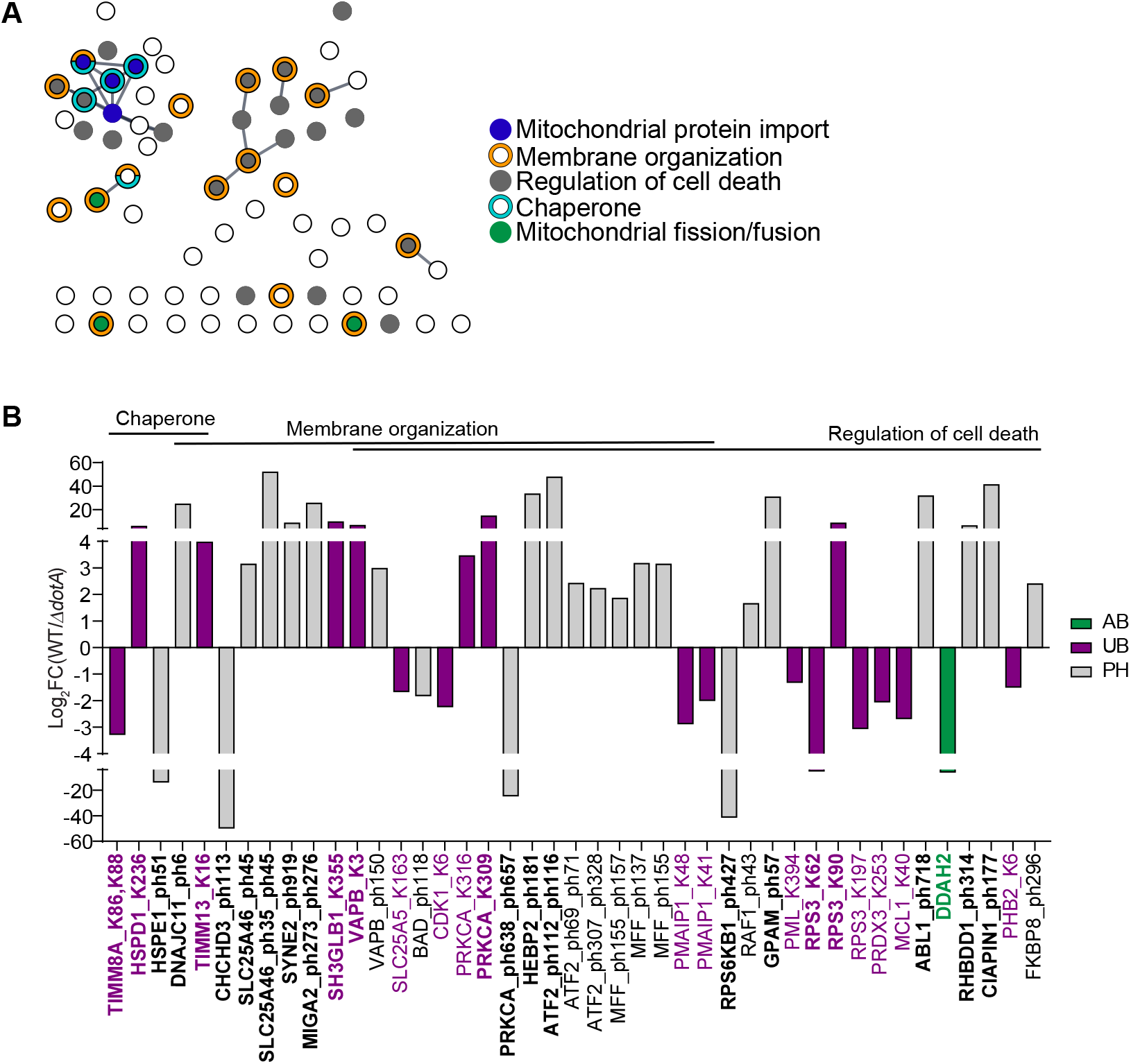
TS44-dependent changes in the mitochondrial proteome in response to *L.p.* infection. Related to Figure 4. (A) Gene ontology enrichment and network analysis of all regulated mitochondrial proteins in WT-*vs. ΔdotA*-infected cells at 1h p.i. (AB, UB, PH combined) was performed with the Cytoscape stringApp (Doncheva et al., 2019; Shannon et al., 2003). Each circle represents one protein. Selected overrepresented pathways are highlighted and annotated. (B) Log_2_FC(WT/ΔdotA) values of significantly regulated mitochondrial proteins/proteoforms from selected gene ontology terms in (A). Shown are changes in protein abundance (green), ubiquitination sites (purple) and phosphorylation sites (grey) at 1h p.i.. Proteins/proteoforms with imputed values are highlighted in bold.

**Figure S5.**
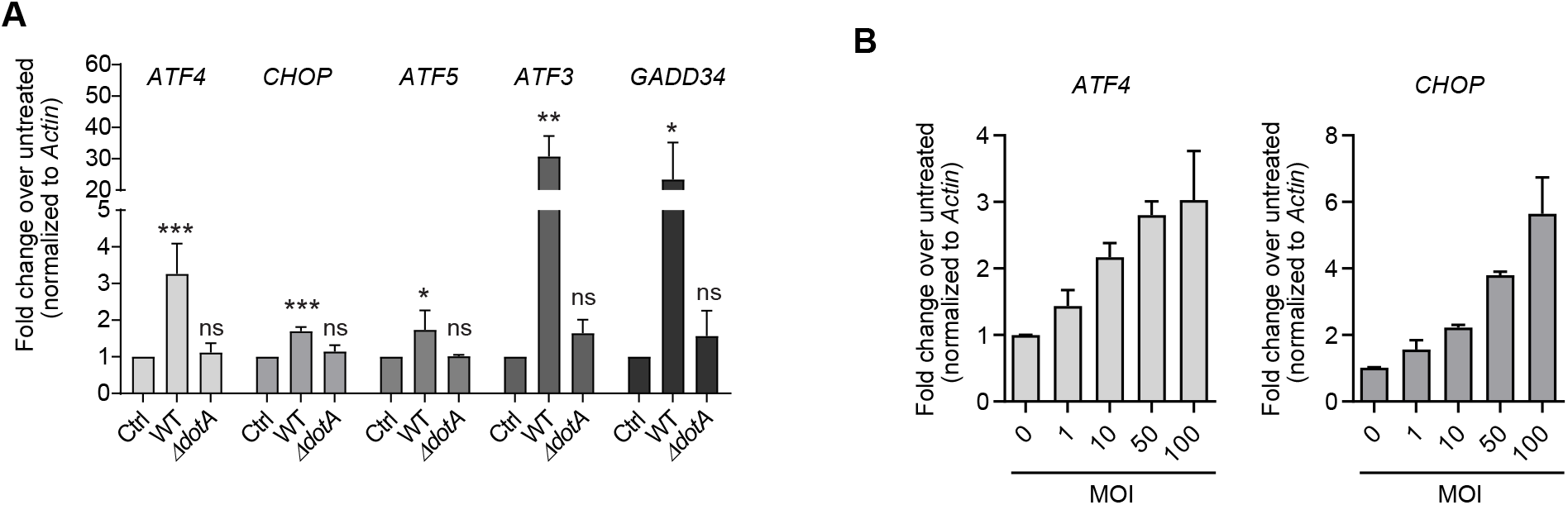
Induction of mitochondrial stress markers during *L.p.* infection. Related to Figure 5. (A) qPCR of *ATF4, CHOP,* ATF5, *ATF3* and *GADD34* in uninfected, WT- or *ΔdotA-infected* HEK293 FcγR (6h p.i., MOI 10). Transcript levels were normalized to *Actin.* Shown are the mean levels relative to the control ± SEM of n = 2 biological replicates. Statistical differences were analyzed by one-way ANOVA and Tukey’s multiple comparison test. *** p ≤ 0.001, ** p ≤ 0.01, * p ≤ 0.05, ns > 0.05. (B) HEK293 FcγR cells were infected with the indicated MOIs for 6h and transcript levels of *ATF4* and *CHOP* were analyzed by qPCR. Shown are the mean levels relative to the control ± SEM of n = 2-3 biological replicates.

**Figure S6.**
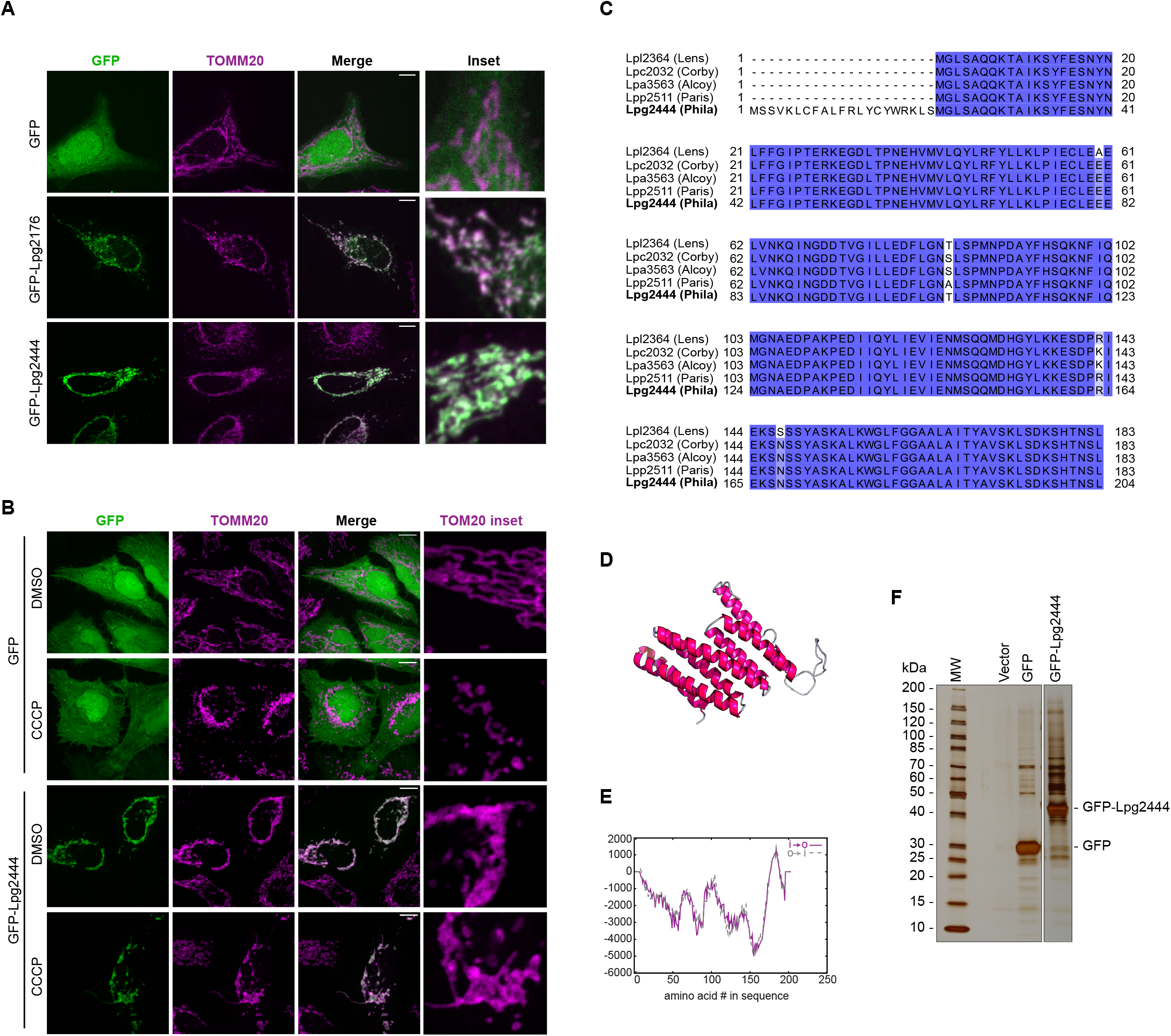
Characterization of the *L.p.* effector Lpg2444. Related to Figure 6. (A) Immunofluorescence analysis of HeLa cells transfected with GFP, GFP-Lpg176 or GFP-Lpg2444. Mitochondria were stained with an anti-TOMM20 antibody (magenta). Scale bar: 10 μm. (B) Immunofluorescence analysis of mitochondrial morphology in GFP-transfected or GFP-Lpg2444-transfected HeLa cells treated with DMSO or 10 μM CCCP for 6h. Mitochondria were stained with an anti-TOM2O antibody (magenta). Scale bar: 10 μm. (C) Amino acid sequence alignment of Lpg2444 *(L.p.* strain Philadelphia) and its homologs Lpl2364 *(L.p.* strain Lens), Lpc2032 *(L.p.* strain Corby) and Lpp2511 (*L.p.* strain Paris). Sequence similarity is indicated by shades of blue. (D) Predicted structure of Lpg2444 using RaptorX (Källberg et al., 2012) using 5cwmA as a template (p-value 2.37e-03). (E) Prediction of transmembrane domains in Lpg2444 with TMPred (Hofmann and Stoffel, 1993). (F) Cell lysates expressing an empty vector control, GFP or GFP-Lpg2444 were subjected to a GFP-Trap affinity resin purification and analyzed by AP-MS. A small aliquot was loaded on a polyacrylamide-gel and proteins were visualized with silver stain.

**Table S3.** Numbers of significantly regulated proteins and proteoforms in uninfected (Ctrl), WT-or *ΔdotA*-infected cells (adj. p-value ≤ 0.05, |Log_2_FC| ≥ 1 in at least 2 out of 3 biological replicates).

| Dataset | Time<br>*p.i.* | WT/Ctrl |  | WT/*ΔdotA* |  | *ΔdotA*/Ctrl |  |
| --- | --- | --- | --- | --- | --- | --- | --- |
|  |  | Proteins<br>(proteoforms) | Proteins<br>(proteoforms) | Proteins<br>(proteoforms) | Proteins<br>(proteoforms) | Proteins<br>(proteoforms) | Proteins<br>(proteoforms) |
| | | $\text{Log}_2\text{FC} \geq 1$<br>adj. $p \leq 0.05$ | $\text{Log}_2\text{FC} \leq -1$<br>adj. $p \leq 0.05$ | $\text{Log}_2\text{FC} \geq 1$<br>adj. $p \leq 0.05$ | $\text{Log}_2\text{FC} \leq -1$<br>adj. $p \leq 0.05$ | $\text{Log}_2\text{FC} \geq 1$<br>adj. $p \leq 0.05$ | $\text{Log}_2\text{FC} \leq -1$<br>adj. $p \leq 0.05$ |
| AB | 1h | 49 | 202 | 38 | 19 | 22 | 233 |
|  | 8h | 103 | 261 | 68 | 227 | 45 | 27 |
| UB | 1h | 420 | 195 | 400 | 76 | 30 | 52 |
|  |  | (868) | (416) | (855) | (225) | (76) | (165) |
|  | 8h | 271 | 189 | 221 | 224 | 62 | 22 |
|  |  | (532) | (381) | (473) | (447) | (141) | (58) |
| PH | 1h | 213 | 264 | 145 | 40 | 51 | 216 |
|  |  | (971) | (987) | (649) | (248) | (260) | (574) |
|  | 8h | 329 | 484 | 322 | 424 | 28 | 28 |
|  |  | (1333) | (2106) | (1301) | (1929) | (153) | (166) |

## REFERENCES

Ahola, S., Langer, T., and MacVicar, T. (2019). Mitochondrial Proteolysis and Metabolic Control. Cold Spring Harb. Perspect. Biol. 11, a033936.

Alkema, M.J., Bronk, M., Verhoeven, E., Otte, A., Van ‘T Veer, L.J., Berns, A., and Van Lohuizen, M. (1997). Identification of Bmi1-interacting proteins as constituents of a multimeric mammalian Polycomb complex. Genes Dev.

Almagro Armenteros, J.J., Sønderby, C.K., Sønderby, S.K., Nielsen, H., and Winther, O. (2017). DeepLoc: prediction of protein subcellular localization using deep learning. Bioinformatics 33, 3387–3395.

An, H., Ordureau, A., Paulo, J.A., Shoemaker, C.J., Denic, V., and Harper, J.W. (2019). TEX264 Is an Endoplasmic Reticulum-Resident ATG8-Interacting Protein Critical for ER Remodeling during Nutrient Stress. Mol. Cell 74, 891–908.e10.

Andréasson, C., Ott, M., and Büttner, S. (2019). Mitochondria orchestrate proteostatic and metabolic stress responses. EMBO Rep. 20, e47865.

Arasaki, K., and Roy, C.R. (2010). Legionella pneumophila promotes functional interactions between plasma membrane syntaxins and Sec22b. Traffic 11, 587–600.

Arasaki, K., Mikami, Y., Shames, S.R., Inoue, H., Wakana, Y., and Tagaya, M. (2017). Legionella effector Lpg1137 shuts down ER-mitochondria communication through cleavage of syntaxin 17. Nat. Commun. 8, 15406.

Banga, S., Gao, P., Shen, X., Fiscus, V., Zong, W.-X., Chen, L., and Luo, Z.-Q. (2007). Legionella pneumophila inhibits macrophage apoptosis by targeting pro-death members of the Bcl2 protein family. Proc. Natl. Acad. Sci. 104, 5121–5126.

Belyi, Y., Niggeweg, R., Opitz, B., Vogelsgesang, M., Hippenstiel, S., Wilm, M., and Aktories, K. (2006). Legionella pneumophila glucosyltransferase inhibits host elongation factor {1A}. Proc. Natl. Acad. Sci. U. S. A. 103, 16953–16958.

Belyi, Y., Tabakova, I., Stahl, M., and Aktories, K. (2008). Lgt: a family of cytotoxic glucosyltransferases produced by Legionella pneumophila. J. Bacteriol. 190, 3026–3035.

Bhogaraju, S., Kalayil, S., Liu, Y., Bonn, F., Colby, T., Matic, I., and Dikic, I. (2016). Phosphoribosylation of Ubiquitin Promotes Serine Ubiquitination and Impairs Conventional Ubiquitination. Cell 167, 1636–1649.e13.

Bhogaraju, S., Bonn, F., Mukherjee, R., Adams, M., Pfleiderer, M.M., Galej, W.P., Matkovic, V., Lopez-Mosqueda, J., Kalayil, S., Shin, D., et al. (2019). Inhibition of bacterial ubiquitin ligases by SidJ-calmodulin catalysed glutamylation. Nature 572, 382–386.

Bruckert, W.M., and Abu Kwaik, Y. (2015). Complete and ubiquitinated proteome of the Legionella-containing vacuole within human macrophages. J. Proteome Res.

Bueno, M., Brands, J., Voltz, L., Fiedler, K., Mays, B., St. Croix, C., Sembrat, J., Mallampalli, R.K., Rojas, M., and Mora, A.L. (2018). ATF3 represses PINK1 gene transcription in lung epithelial cells to control mitochondrial homeostasis. Aging Cell 17, e12720.

Carpenter, A.E., Jones, T.R., Lamprecht, M.R., Clarke, C., Kang, I., Friman, O., Guertin, D.A., Chang, J., Lindquist, R.A., Moffat, J., et al. (2006). CellProfiler: image analysis software for identifying and quantifying cell phenotypes. Genome Biol. 7, R100.

Chino, H., Hatta, T., Natsume, T., and Mizushima, N. (2019). Intrinsically Disordered Protein TEX264 Mediates ER-phagy. Mol. Cell 74, 909–921.e6.

Choi, M., Chang, C.Y., Clough, T., Broudy, D., Killeen, T., MacLean, B., and Vitek, O. (2014). MSstats: An R package for statistical analysis of quantitative mass spectrometry-based proteomic experiments. Bioinformatics 30, 2524–2526.

Claros, M.G., and Vincens, P. (1996). Computational method to predict mitochondrially imported proteins and their targeting sequences. Eur. J. Biochem. 241, 779–786.

Cornejo, E., Schlaermann, P., and Mukherjee, S. (2017). How to rewire the host cell: A home improvement guide for intracellular bacteria. J. Cell Biol. 216, 3931–3948.

Cox, J., and Mann, M. (2008). MaxQuant enables high peptide identification rates, individualized p.p.b.-range mass accuracies and proteome-wide protein quantification. Nat. Biotechnol. 26, 1367–1372.

D’Amico, D., Sorrentino, V., and Auwerx, J. (2017). Cytosolic Proteostasis Networks of the Mitochondrial Stress Response. Trends Biochem. Sci. 42.

Degtyar, E., Zusman, T., Ehrlich, M., and Segal, G. (2009). A Legionella effector acquired from protozoa is involved in sphingolipids metabolism and is targeted to the host cell mitochondria. Cell. Microbiol. 11, 1219–1235.

Dolezal, P., Aili, M., Tong, J., Jiang, J.-H., Marobbio, C.M., Lee, S. fung, Schuelein, R., Belluzzo, S., Binova, E., Mousnier, A., et al. (2012). Legionella pneumophila Secretes a Mitochondrial Carrier Protein during Infection. PLoS Pathog. 8, e1002459.

Doncheva, N.T., Morris, J.H., Gorodkin, J., and Jensen, L.J. (2019). Cytoscape StringApp: Network Analysis and Visualization of Proteomics Data. J. Proteome Res. 18, 623–632.

Emanuelsson, O., Nielsen, H., Brunak, S., and von Heijne, G. (2000). Predicting subcellular localization of proteins based on their N-terminal amino acid sequence. J. Mol. Biol. 300, 1005–1016.

Emmerich, C.H., and Cohen, P. (2015). Optimising methods for the preservation, capture and identification of ubiquitin chains and ubiquitylated proteins by immunoblotting. Biochem. Biophys. Res. Commun. 466, 1–14.

Escoll, P., Song, O.-R., Viana, F., Steiner, B., Lagache, T., Olivo-Marin, J.-C., Impens, F., Brodin, P., Hilbi, H., and Buchrieser, C. (2017). Legionella pneumophila Modulates Mitochondrial Dynamics to Trigger Metabolic Repurposing of Infected Macrophages. Cell Host Microbe 22, 302–316.e7.

Fessler, E., Eckl, E.-M., Schmitt, S., Mancilla, I.A., Meyer-Bender, M.F., Hanf, M., Philippou-Massier, J., Krebs, S., Zischka, H., and Jae, L.T. (2020). A pathway coordinated by DELE1 relays mitochondrial stress to the cytosol. Nature 579, 433–437.

Finsel, I., and Hilbi, H. (2015). Formation of a pathogen vacuole according to Legionella pneumophila: how to kill one bird with many stones. Cell. Microbiol. 17, 935–950.

Fiorese, C.J., Schulz, A.M., Lin, Y.-F., Rosin, N., Pellegrino, M.W., and Haynes, C.M. (2016). The Transcription Factor ATF5 Mediates a Mammalian Mitochondrial UPR. Curr. Biol. 26, 2037–2043.

Fontana, M.F., Banga, S., Barry, K.C., Shen, X., Tan, Y., Luo, Z.Q., and Vance, R.E. (2011). Secreted bacterial effectors that inhibit host protein synthesis are critical for induction of the innate immune response to virulent Legionella pneumophila. PLoS Pathog.

Fukasawa, Y., Tsuji, J., Fu, S.-C., Tomii, K., Horton, P., and Imai, K. (2015). MitoFates: Improved Prediction of Mitochondrial Targeting Sequences and Their Cleavage Sites. Mol. Cell. Proteomics 4, 1113–1126.

Guo, X., Aviles, G., Liu, Y., Tian, R., Unger, B.A., Lin, Y.-H.T., Wiita, A.P., Xu, K., Correia, M.A., and Kampmann, M. (2020). Mitochondrial stress is relayed to the cytosol by an OMA1-DELE1-HRI pathway. Nature 579, 427–432.

Halestrap, A.P., and Davidson, A.M. (1990). Inhibition of Ca2 -induced large-amplitude swelling of liver and heart mitochondria by cyclosporin is probably caused by the inhibitor binding to mitochondrial-matrix peptidyl-prolyl cis-trans isomerase and preventing it interacting with the adenine nucle. Biochem. J. 268, 153–160.

Hofmann, K., and Stoffel, W. (1993). TMbase: A Database of Membrane Spanning Protein Segments.

Horenkamp, F.A., Mukherjee, S., Alix, E., Schauder, C.M., Hubber, A.M., Roy, C.R., and Reinisch, K.M. (2014). *Legionella pneumophila* Subversion of Host Vesicular Transport by SidC Effector Proteins. Traffic 15, 488–499.

Horwitz, M.A. (1983). Formation of a novel phagosome by the Legionnaires’ disease bacterium (Legionella pneumophila) in human monocytes. J. Exp. Med. 158, 1319–1331.

Hoshino, A., Wang, W.-J., Wada, S., McDermott-Roe, C., Evans, C.S., Gosis, B., Morley, M.P., Rathi, K.S., Li, J., Li, K., et al. (2019). The ADP/ATP translocase drives mitophagy independent of nucleotide exchange. Nature 575, 375–379.

Ishihara, N., Jofuku, A., Eura, Y., and Mihara, K. (2003). Regulation of mitochondrial morphology by membrane potential, and DRP1-dependent division and FZO1-dependent fusion reaction in mammalian cells. Biochem. Biophys. Res. Commun. 301, 891–898.

Ivanov, S.S., and Roy, C.R. (2013). Pathogen signatures activate a ubiquitination pathway that modulates the function of the metabolic checkpoint kinase mTOR. Nat. Immunol. 14, 1219–1228.

Jadhav, K., and Zhang, Y. (2017). Activating transcription factor 3 in immune response and metabolic regulation. Liver Res. 1, 96–102.

Jaeschke, A., Karasarides, M., Ventura, J.-J., Ehrhardt, A., Zhang, C., Flavell, R.A., Shokat, K.M., and Davis, R.J. (2006). JNK2 Is a Positive Regulator of the cJun Transcription Factor. Mol. Cell 23, 899–911.

Jensen, L.J., Kuhn, M., Stark, M., Chaffron, S., Creevey, C., Muller, J., Doerks, T., Julien, P., Roth, A., Simonovic, M., et al. (2009). STRING 8-a global view on proteins and their functional interactions in 630 organisms. Nucleic Acids Res. 37, D412–D416.

Jimenez-Morales, D., Campos, A.R., and Von Dollen, J. (2019). artMS: Analytical R tools for Mass Spectrometry. Bioconductor.

Jovaisaite, V., Mouchiroud, L., and Auwerx, J. (2014). The mitochondrial unfolded protein response, a conserved stress response pathway with implications in health and disease. J. Exp. Biol. 217, 137–143.

Kagan, J.C., Stein, M.-P., Pypaert, M., and Roy, C.R. (2004). Legionella subvert the functions of Rab1 and Sec22b to create a replicative organelle. J. Exp. Med. 199, 1201–1211.

Kalayil, S., Bhogaraju, S., Bonn, F., Shin, D., Liu, Y., Gan, N., Basquin, J., Grumati, P., Luo, Z.-Q., and Dikic, I. (2018). Insights into catalysis and function of phosphoribosyl-linked serine ubiquitination. Nature 557, 734–738.

Källberg, M., Wang, H., Wang, S., Peng, J., Wang, Z., Lu, H., and Xu, J. (2012). Template-based protein structure modeling using the RaptorX web server. Nat. Protoc. 7, 1511–1522.

Kallunki, T., Su, B., Tsigelny, I., Sluss, H.K., Derijard, B., Moore, G., Davis, R., and Karin, M. (1994). JNK2 contains a specificity-determining region responsible for efficient c-Jun binding and phosphorylation. Genes Dev.

Kang, B.H., Plescia, J., Song, H.Y., Meli, M., Colombo, G., Beebe, K., Scroggins, B., Neckers, L., and Altieri, D.C. (2009). Combinatorial drug design targeting multiple cancer signaling networks controlled by mitochondrial Hsp90. J. Clin. Invest. 119, 454–464.

Karch, J., Bround, M.J., Khalil, H., Sargent, M.A., Latchman, N., Terada, N., Peixoto, P.M., and Molkentin, J.D. (2019). Inhibition of mitochondrial permeability transition by deletion of the ANT family and CypD. Sci Adv 5, eaaw4597.

Kenny, T.C., Craig, A.J., Villanueva, A., and Germain, D. (2019). Mitohormesis Primes Tumor Invasion and Metastasis. Cell Rep. 27, 2292–2303.e6.

Kim, W., Bennett, E.J., Huttlin, E.L., Guo, A., Li, J., Possemato, A., Sowa, M.E., Rad, R., Rush, J., Comb, M.J., et al. (2011). Systematic and quantitative assessment of the ubiquitin-modified proteome. Mol. Cell 44, 325–340.

Kotewicz, K.M., Ramabhadran, V., Sjoblom, N., Vogel, J.P., Haenssler, E., Zhang, M., Behringer, J., Scheck, R.A., and Isberg, R.R. (2017). A Single Legionella Effector Catalyzes a Multistep Ubiquitination Pathway to Rearrange Tubular Endoplasmic Reticulum for Replication. Cell Host Microbe 21, 169–181.

Laguna, R.K., Creasey, E.A., Li, Z., Valtz, N., and Isberg, R.R. (2006). A Legionella pneumophila-translocated substrate that is required for growth within macrophages and protection from host cell death. Proc. Natl. Acad. Sci. U. S. A. 103, 18745–18750.

Lebeau, J., Rainbolt, T.K., and Wiseman, R.L. (2018). Coordinating Mitochondrial Biology Through the Stress-Responsive Regulation of Mitochondrial Proteases. Int. Rev. Cell Mol. Biol. 340, 79–128.

Li, T., Lu, Q., Wang, G., Xu, H., Huang, H., Cai, T., Kan, B., Ge, J., and Shao, F. (2013). SET-domain bacterial effectors target heterochromatin protein 1 to activate host rDNA transcription. EMBO Rep. 14, 733–740.

López-Sánchez, I., Sanz-Garcia, M., and Lazo, P.A. (2009). Plk3 interacts with and specifically phosphorylates VRK1 in Ser342, a downstream target in a pathway that induces Golgi fragmentation. Mol. Cell. Biol. 29, 1189–1201.

MacVicar, T., Ohba, Y., Nolte, H., Mayer, F.C., Tatsuta, T., Sprenger, H.-G., Lindner, B., Zhao, Y., Li, J., Bruns, C., et al. (2019). Lipid signalling drives proteolytic rewiring of mitochondria by YME1L. Nature 575, 361–365.

Matic, S., Muders, V., and Meisinger, C. (2018). Tuning the mitochondrial protein import machinery by reversible phosphorylation: from metabolic switches to cell cycle regulation. Curr. Opin. Physiol. 3.

McLelland, G.-L., Goiran, T., Yi, W., Dorval, G., Chen, C.X., Lauinger, N.D., Krahn, A.I., Valimehr, S., Rakovic, A., Rouiller, I., et al. (2018). Mfn2 ubiquitination by PINK1/parkin gates the p97-dependent release of ER from mitochondria to drive mitophagy. Elife 7, e32866.

Metz, K.S., Deoudes, E.M., Berginski, M.E., Jimenez-Ruiz, I., Aksoy, B.A., Hammerbacher, J., Gomez, S.M., and Phanstiel, D.H. (2018). Coral: Clear and Customizable Visualization of Human Kinome Data. Cell Syst. 7, 347–350.e1.

Michard, C., and Doublet, P. (2015). Post-translational modifications are key players of the Legionella pneumophila infection strategy. Front. Microbiol. 6, 87.

Moss, S.M., Taylor, I.R., Ruggero, D., Gestwicki, J.E., Shokat, K.M., and Mukherjee, S. (2019). A Legionella pneumophila Kinase Phosphorylates the Hsp70 Chaperone Family to Inhibit Eukaryotic Protein Synthesis. Cell Host Microbe 25, 454–462.e6.

Mottis, A., Herzig, S., and Auwerx, J. (2019). Mitocellular communication: Shaping health and disease. Science (80-.). 366, 827–832.

Mukherjee, S., Liu, X., Arasaki, K., McDonough, J., Galán, J.E., and Roy, C.R. (2011). Modulation of Rab GTPase function by a protein phosphocholine transferase. Nature 477, 103–106.

Münch, C., and Harper, J.W. (2016). Mitochondrial unfolded protein response controls matrix pre-RNA processing and translation. Nature 534, 710–713.

Noack, J., and Mukherjee, S. (2020). “Make way”: Pathogen exploitation of membrane traffic. Curr. Opin. Cell Biol.

Ochoa, D., Jonikas, M., Lawrence, R.T., El Debs, B., Selkrig, J., Typas, A., Villén, J., Santos, S.D., and Beltrao, P. (2016). An atlas of human kinase regulation. Mol. Syst. Biol. 12, 888.

Pakos-Zebrucka, K., Koryga, I., Mnich, K., Ljujic, M., Samali, A., and Gorman, A.M. (2016). The integrated stress response. EMBO Rep. 17, 1374–1395.

Pellegrino, M.W., Nargund, A.M., Kirienko, N. V., Gillis, R., Fiorese, C.J., and Haynes, C.M. (2014). Mitochondrial UPR-regulated innate immunity provides resistance to pathogen infection. Nature 516, 414–417.

Peng, H., Yang, J., Li, G., You, Q., Han, W., Li, T., Gao, D., Xie, X., Lee, B.H., Du, J., et al. (2017). Ubiquitylation of p62/sequestosome1 activates its autophagy receptor function and controls selective autophagy upon ubiquitin stress. Cell Res. 27, 657–674.

Qiu, J., and Luo, Z.-Q. (2017a). Legionella and Coxiella effectors: strength in diversity and activity. Nat. Rev. Microbiol. 15, 591–605.

Qiu, J., and Luo, Z.-Q. (2017b). Hijacking of the Host Ubiquitin Network by Legionella pneumophila. Front. Cell. Infect. Microbiol. 7, 487.

Qiu, J., Sheedlo, M.J., Yu, K., Tan, Y., Nakayasu, E.S., Das, C., Liu, X., and Luo, Z.-Q. (2016). Ubiquitination independent of E1 and E2 enzymes by bacterial effectors. Nature 533, 120–124.

Quirós, P.M., Prado, M.A., Zamboni, N., D’Amico, D., Williams, R.W., Finley, D., Gygi, S.P., and Auwerx, J. (2017). Multi-omics analysis identifies ATF4 as a key regulator of the mitochondrial stress response in mammals. J. Cell Biol. 216, 2027–2045.

Rainbolt, T.K., Atanassova, N., Genereux, J.C., and Wiseman, R.L. (2013). Stress-regulated translational attenuation adapts mitochondrial protein import through Tim17A degradation. Cell Metab. 18, 908–919.

Raudvere, U., Kolberg, L., Kuzmin, I., Arak, T., Adler, P., Peterson, H., and Vilo, J. (2019). g:Profiler: a web server for functional enrichment analysis and conversions of gene lists (2019 update). Nucleic Acids Res. 47, W191–W198.

Rolando, M., Sanulli, S., Rusniok, C., Gomez-Valero, L., Bertholet, C., Sahr, T., Margueron, R., and Buchrieser, C. (2013). Legionella pneumophila effector RomA uniquely modifies host chromatin to repress gene expression and promote intracellular bacterial replication. Cell Host Microbe.

Schmidt, O., Harbauer, A.B., Rao, S., Eyrich, B., Zahedi, R.P., Stojanovski, D., Schönfisch, B., Guiard, B., Sickmann, A., Pfanner, N., et al. (2011). Regulation of mitochondrial protein import by cytosolic kinases. Cell 144, 227–239.

Schmölders, J., Manske, C., Otto, A., Hoffmann, C., and others (2017). Comparative proteomics of purified pathogen vacuoles correlates intracellular replication of Legionella pneumophila with the small GTPase ras-related protein 1 (Rap1). Mol. Cell. Proteomics 16, 622–641.

Shannon, P., Markiel, A., Ozier, O., Baliga, N.S., Wang, J.T., Ramage, D., Amin, N., Schwikowski, B., and Ideker, T. (2003). Cytoscape: a software environment for integrated models of biomolecular interaction networks. Genome Res. 13, 2498–2504.

Shen, X., Banga, S., Liu, Y., Xu, L., Gao, P., Shamovsky, I., Nudler, E., and Luo, Z.-Q. (2009). Targeting eEF1A by a Legionella pneumophila effector leads to inhibition of protein synthesis and induction of host stress response. Cell. Microbiol. 11, 911–926.

Shin, D., Mukherjee, R., Liu, Y., Gonzalez, A., Bonn, F., Liu, Y., Rogov, V. V, Heinz, M., Stolz, A., Hummer, G., et al. (2020). Regulation of Phosphoribosyl-Linked Serine Ubiquitination by Deubiquitinases DupA and DupB. Mol. Cell 77, 164–179.e6.

Shpilka, T., and Haynes, C.M. (2017). The mitochondrial UPR: mechanisms, physiological functions and implications in ageing. Nat. Rev. Mol. Cell Biol. 19, 109–120.

Sidrauski, C., Acosta-Alvear, D., Khoutorsky, A., Vedantham, P., Hearn, B.R., Li, H., Gamache, K., Gallagher, C.M., Ang, K.K.-H., Wilson, C., et al. (2013). Pharmacological brake-release of mRNA translation enhances cognitive memory. Elife 2, e00498.

Siegelin, M.D., Dohi, T., Raskett, C.M., Orlowski, G.M., Powers, C.M., Gilbert, C.A., Ross, A.H., Plescia, J., and Altieri, D.C. (2011). Exploiting the mitochondrial unfolded protein response for cancer therapy in mice and human cells. J. Clin. Invest. 121, 1349–1360.

Sun, S., and Habermann, B.H. (2017). A Guide to Computational Methods for Predicting Mitochondrial Localization. In Mitochondria: Practical Protocols, D. Mokranjac, and F. Perocchi, eds. (New York, NY: Springer New York), pp. 1–14.

Swaney, D.L., and Villén, J. (2016). Enrichment of Phosphopeptides via Immobilized Metal Affinity Chromatography. Cold Spring Harb. Protoc. 2016, db.prot088005.

Tanaka, A., Cleland, M.M., Xu, S., Narendra, D.P., Suen, D.-F., Karbowski, M., and Youle, R.J. (2010). Proteasome and p97 mediate mitophagy and degradation of mitofusins induced by Parkin. J. Cell Biol. 191, 1367–1380.

Teo, G., Liu, G., Zhang, J., Nesvizhskii, A.I., Gingras, A.C., and Choi, H. (2014). SAINTexpress: Improvements and additional features in Significance Analysis of INTeractome software. J. Proteomics 100, 37–43.

Thul, P.J., Akesson, L., Wiking, M., Mahdessian, D., Geladaki, A., Ait Blal, H., Alm, T., Asplund, A., Björk, L., Breckels, L.M., et al. (2017). A subcellular map of the human proteome. Science (80-.). 356, eaal3321.

Tiku, V., Kew, C., Mehrotra, P., Ganesan, R., Robinson, N., and Antebi, A. (2018). Nucleolar fibrillarin is an evolutionarily conserved regulator of bacterial pathogen resistance. Nat. Commun. 9.

Tiku, V., Tan, M.-W., and Dikic, I. (2020). Mitochondrial Functions in Infection and Immunity. Trends Cell Biol. 30, 263–275.

Treacy-Abarca, S., and Mukherjee, S. (2015). Legionella suppresses the host unfolded protein response via multiple mechanisms. Nat. Commun. 6, 7887.

Udeshi, N.D., Mani, D.R., Eisenhaure, T., Mertins, P., Jaffe, J.D., Clauser, K.R., Hacohen, N., and Carr, S.A. (2012). Methods for quantification of in vivo changes in protein ubiquitination following proteasome and deubiquitinase inhibition. Mol. Cell. Proteomics 11, 148–159.

Urwyler, S., Nyfeler, Y., Ragaz, C., Lee, H., Mueller, L.N., Aebersold, R., and Hilbi, H. (2009). Proteome analysis of Legionella vacuoles purified by magnetic immunoseparation reveals secretory and endosomal GTPases. Traffic 10, 76–87.

Vizcaíno, J.A., Csordas, A., del-Toro, N., Dianes, J.A., Griss, J., Lavidas, I., Mayer, G., Perez-Riverol, Y., Reisinger, F., Ternent, T., et al. (2016). 2016 update of the PRIDE database and its related tools. Nucleic Acids Res. 44, D447–56.

Wang, Y., Wan, B., Li, D., Zhou, J., Li, R., Bai, M., Chen, F., and Yu, L. (2012). BRSK2 is regulated by ER stress in protein level and involved in ER stress-induced apoptosis. Biochem. Biophys. Res. Commun. 423, 813–818.

Wang, Y., Shi, M., Feng, H., Zhu, Y., Liu, S., Gao, A., and Gao, P. (2018). Structural Insights into Non-canonical Ubiquitination Catalyzed by SidE. Cell 173, 1231–1243.e16.

Webb-Robertson, B.-J.M., Wiberg, H.K., Matzke, M.M., Brown, J.N., Wang, J., McDermott, J.E., Smith, R.D., Rodland, K.D., Metz, T.O., Pounds, J.G., et al. (2015). Review, evaluation, and discussion of the challenges of missing value imputation for mass spectrometry-based label-free global proteomics. J. Proteome Res. 14, 1993–2001.

Yu, C.-S., Chen, Y.-C., Lu, C.-H., and Hwang, J.-K. (2006). Prediction of protein subcellular localization. Proteins 64, 643–651.

Zhu, B., Zheng, Y., Pham, A.D., Mandal, S.S., Erdjument-Bromage, H., Tempst, P., and Reinberg, D. (2005). Monoubiquitination of human histone H2B: The factors involved and their roles in HOX gene regulation. Mol. Cell 20, 601–611.

